# Derivation of macaque trophoblast stem cells

**DOI:** 10.1101/2020.03.17.995407

**Authors:** Jenna Kropp Schmidt, Michael G. Meyer, Gregory J. Wiepz, Lindsey N. Block, Brittany M. Dusek, Kamryn M. Kroner, Mario J. Bertogliat, Logan T. Keding, Michelle R. Koenig, Katherine D. Mean, Thaddeus G. Golos

**Affiliations:** Wisconsin National Primate Research Center, University of Wisconsin-Madison; Department of Comparative Biosciences, University of Wisconsin-Madison; Department of Obstetrics and Gynecology, University of Wisconsin-Madison

**Author notes:** Corresponding Author: Jenna Kropp Schmidt, Wisconsin National Primate Research Center, 1220 Capitol Ct., Madison, WI 53965. Competing Interests: Authors declare no competing interests.

**Keywords:** trophoblast, macaque, stem cell, placenta

## Abstract

Nonhuman primates are excellent models for studying human placentation as experimental manipulations *in vitro* can be translated to *in vivo* pregnancy. Our objective was to develop macaque trophoblast stem cells (TSC) as an *in vitro* platform for future assessment of primate trophoblast development and function. Macaque TSC lines were generated by isolating first trimester placental villous cytotrophoblasts followed by culture in TSC medium to “reprogram” the cells to a proliferative state. TSCs grew as mononuclear colonies, whereas upon induction of syncytiotrophoblast (ST) differentiation multinuclear structures appeared, indicative of syncytium formation. Chorionic gonadotropin secretion was >4,000-fold higher in ST culture media compared to TSC media. Characteristic trophoblast hallmarks were defined in TSCs and ST including expression of C19MC miRNAs and macaque placental nonclassical MHC class I molecule, Mamu-AG. TSC differentiation to extravillous trophoblasts (EVTs) with or without the ALK-5 inhibitor A83-01 resulted in differing morphologies but similar expression of Mamu-AG and CD56 as assessed by flow cytometry, hence further refinement of relevant EVT markers is needed. Our preliminary characterization of macaque TSCs suggests that these cells represent a proliferative, self-renewing TSC population capable of differentiating to STs *in vitro* thereby establishing an experimental model of primate placentation.

## Introduction

Pregnancy-related complications, including preeclampsia and intrauterine fetal growth restriction, stem from suboptimal placental development^1, 2^. The consequences of poor placental development transcend pregnancy as fetal programming in utero predisposes the fetus to increased risk for cardiovascular disease and metabolic syndrome in adulthood^3, 4^. The molecular cues underlying placental developmental events occurring within the first trimester of human pregnancy, however, are not well understood. Therefore, defining the mechanisms underlying early placentation is necessary to not only improve both maternal and fetal well-being, but also to improve the child’s health throughout their lifespan.

Early placental development in humans is relatively unexplored as *in vivo* samples are a limited resource. Moreover, there is restricted ability to functionally perturb gene networks or evaluate cell-type specific responses to pathogens or other environmental factors at the maternal-fetal interface in human tissue. In the mouse, trophoblast stem cells (TSCs) can be derived by culturing extraembryonic ectoderm in the presence of FGF4, heparin and mouse embryonic fibroblasts (or supplementation of TGF-ß1/Activin) and can readily serve as an *in vitro* experimental platform^5^. Notably, the timing of developmental events^6, 7^, organization of cell types^6, 8^ and trophoblast gene expression profiles^6, 9–11^ differ greatly between the mouse and human. In addition, the conditions that drive mouse TSC derivation fail to support the differentiation of human TSCs^12^. These differences necessitate the use of human or primate embryos, trophoblast cell lines, and placental tissue for elucidating early developmental circuitry.

Human trophoblast cell models include cells derived from naturally occurring choriocarcinomas (AC1M, BeWo, JAR, JEG-3)^13–15^, immortalization of human primary trophoblast cultures (HTR-8/SVneo, SWAN-71)^13, 16, 17^, treatment of human pluripotent embryonic stem cells (ESCs) to promote specification of trophectoderm^18, 19^, and the isolation of primary trophoblasts for in vitro culture^20^. Human ESCs upon treatment with BMP differentiate to trophoblast-like cells^18, 19^, however, these cells do not continue to proliferate nor is differentiation tightly controlled. There is also considerable variation in BMP4 treatment protocols leading to differing expression profiles and phenotypes^21, 22^. The ESC derived trophoblasts express gene signatures representative of a more primitive trophoblast cell that would be present near the time of implantation^21^, as their gene expression profiles differ from primary villous cytotrophoblasts. An alternative approach to obtaining trophoblast cell lines has been to isolate primary trophoblasts from early first trimester placental tissue^22^. It is postulated that a TSC niche resides in the early human first trimester placenta^8, 22–24^, where these cells are a proliferative, less differentiated progenitor cell population that gives rise to both villous cytotrophoblasts that fuse to form the syncytia as well as extravillous trophoblast (EVT) progenitor cells at the cell column tips^25^. Genbacev et al.^26^ isolated primary human trophoblast progenitor cells from the chorion. While these cells demonstrated both trophoblast (cytokeratin-7) and pluripotency (OCT4) marker expression, cell proliferation could not be maintained without differentiation beyond passages 8-10. James et al.^27^ isolated a trophoblast side-population following enzymatic digestion of first trimester villous tissue by sorting cells that weakly or negatively stain for Hoescht 33342^27^, where efflux of the stain is a characteristic of adult stem cells. Self-renewal of the trophoblast side-population was not demonstrated nor have the cells been characterized in their ability to differentiate to both syncytiotrophoblasts (ST) or EVTs. Altogether, the isolation and characterization of human TSC populations has been limited given the uncertainty of which markers truly define primitive or early trophoblasts and TSCs^22^.

Recent optimization of culture conditions for primary human first trimester trophoblasts has led to the development of *in vitro* trophoblast stem cell lines^28–30^. Okae et al.^29^ defined a TSC medium capable of supporting continued proliferation of primary human cytotrophoblasts and blastocyst-derived trophoblasts when cultured on a collagen type IV matrix. TSCs displayed trophoblast transcription factor expression including TEAD4, TP63, and GATA3, as well as expression of chromosome 19 microRNA cluster (C19MC) miRNAs and DNA methylation signatures characteristic of trophoblasts. Importantly, under differentiation-specific culture media formulations TSCs could be differentiated to either chorionic gonadotropin (CG)-secreting syncytia or HLA-G positive extravillous trophoblasts (EVTs). In a different approach, Haider et al.^28^ established a three-dimensional trophoblast organoid model by embedding human cytotrophoblasts, isolated from 6-7 weeks of gestation, into Matrigel and culturing cells in defined medium. The organoid structures formed mononuclear trophoblasts on the outer periphery with an inner layer of CGß expressing multinuclear syncytia. The organoids were maintained for up to five passages and upon removal of R-spondin and CHIR99021 from the culture media, trophoblastic outgrowths formed that were HLA-G positive, a hallmark of human EVTs. A trophoblast organoid model was also developed by Turco et al.^30^, where following enzymatic digestion of 6-9 week human placental tissue, cell clusters were embedded into Matrigel. Similar to the trophoblast organoids generated by Haider et al.^28^, these organoids formed mononuclear cells on the periphery with syncytia and lacuna structures within the center. The trophoblast organoids could be maintained long-term and were also capable of differentiating to EVTs. Although the human TSC media components vary across these studies, collectively they have demonstrated that Wnt activation, EGF signaling, and inhibition of TGF-ß are essential to the maintenance of primary human trophoblasts in culture.

Macaques are an ideal model for human pregnancy studies as, like the human, they develop a villous hemochorial placenta. Primate placentation is characterized by invasion of trophoblasts into the decidualized endometrium and remodeling of maternal spiral arteries. Importantly, macaques express primate and placenta-specific MHC class I homologs^31–34^ and C19MC miRNAs^35^ similar to humans. The macaque model also allows for the study of placentation in its entirety from embryonic trophoblast specification to placental development *in vivo*. Macaque *in vitro* models have been restricted to extended culture of hatched blastocysts ^36–38^, which are not easily controlled. For instance, the derivation of rhesus macaque TSCs was previously reported by VandeVoort et al.^38^ by extended culture of rhesus macaque blastocysts. The trophoblast outgrowths were capable of differentiation as evident by upregulation of CG protein expression upon treatment with 17ß-estradiol. The blastocyst-derived TSCs were capable of being passaged, however, this model and the rhesus macaque *in vitro* embryo outgrowth models^36, 37^ lack the ability to tightly control differentiation in a cell-type specific manner.

The NHP model presents an experimental continuum utilizing *in vitro* embryos, *in vitro* trophoblast cell cultures and experimental *in vivo* pregnancy studies to encompass each stage of pregnancy. Thus, our objective was to derive macaque TSCs utilizing the method described by Okae et al.^29^ to generate human TSCs. We hypothesized that rhesus macaque placental cytotrophoblasts could be “reprogrammed” to a proliferative state upon culture in TSC medium optimized for primate TSCs. Macaque TSC lines were generated by isolating first trimester macaque placental villous cytotrophoblasts followed by culture in TSC medium optimized for human trophoblasts. Here we demonstrate that these trophoblasts can be cultured long-term, display trophoblast marker criteria described by Lee et al.^39^, and are capable of differentiation to both ST and EVT cells. The macaque TS model offers an experimental platform for developing and optimizing genetic tools to assess trophoblast function *in vitro* for functional translation to *in vivo* macaque pregnancy studies.

## Results

### Generation of TSCs and Differentiated Trophoblast Cells

Placentas were collected between 40-75 days of gestation from eight pregnant macaques to isolate primary villous cytotrophoblasts (pri-CTB) for reprogramming to self-renewing TSCs, as illustrated in Figure 1A. A total of three male and three female rhesus, and one each male and female cynomolgus TSC lines were derived (Figure 1B). Pri-CTB cultured in non-TSC media for 72 h readily formed syncytia, while TSC cultures initially displayed mixed cellular morphologies. (Figure 1C, Supplemental Figure S1). Upon extended culture of pri-CTB in TSC medium, TSC cultures predominantly presented as mononuclear cell colonies (Figure 1D). Normal karyotypes were observed in four lines, while an abnormal karyotype was observed in the other four lines with a subset of cells being polyploid (Supplemental Table S1). Of note, chromosome integrity may drift as the cells are passaged, as rh052318 TSCs displayed an abnormal karyotype at a later passage, while analysis of an earlier cryopreserved passage confirmed a normal karyotype. TSC lines have been cultured up to 50 passages with consistent mononuclear TSC-like morphologies, but comprehensive characterization of karyotype and marker expression of later passages has not yet been performed.

**Figure 1.**
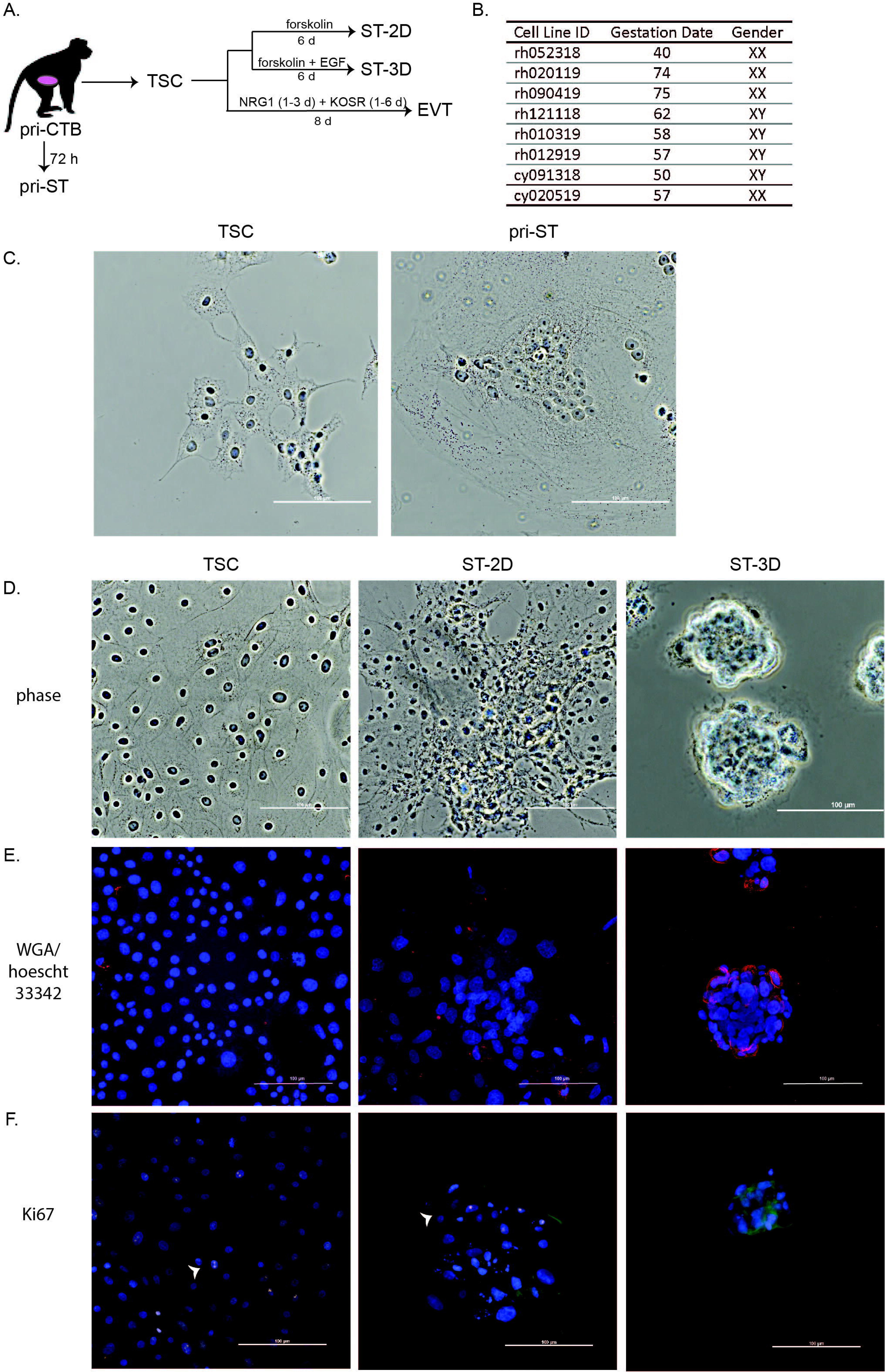
Generation of trophoblast stem cells (TSCs) and differentiated trophoblasts cell types. A) Primary villous cytotrophoblasts (pri-CTB) were isolated from macaque placentas and differentiated to primary syncytiotrophoblasts (pri-ST) or reprogrammed to TSCs for subsequent differentiation. Treatment for directing TSCs to adherent 2-dimensional syncytiotrophoblasts (ST-2D), 3-dimensional syncytiotrophoblasts (ST-3D) cultured in suspension, or adherent extravillous cytotrophoblasts (EVT) are outlined. B) Summary of eight TSC lines derived from first or early second trimester rhesus (rh) or cynomolgus (cy) macaque placentas. C) Representative phase-contrast images of TSC and pri-ST at 72 h of culture. D) Representative phase-contrast microscopic images of each cell type. E) Cells stained with wheat-germ agglutinin (red) to display cell membranes and Hoescht 33342 to visualize nuclei. F) Cell proliferation depicted by Ki-67 immunostaining (punctate pink staining, white arrowhead), f-actin lectin staining (green), and nuclei stained by DAPI (blue). Scale bars in all panels represent 100 µm.

Putative TSCs were differentiated to syncytiotrophoblasts in 2-dimensional (ST-2D) and 3-dimensional (ST-3D) paradigms by either plating TSCs as adherent cultures on collagen IV (col IV)-coated dishes or by culturing in suspension, respectively (Figure 1A). Adherent ST-2D cells displayed a more “raised” structure compared to TSCs, with the presence of both multinuclear and mononuclear cell regions indicative of inconsistent syncytialization. TSCs cultured in suspension with ST medium (ST-3D), largely formed multinuclear aggregates that increased in size as cells aggregated throughout the 6 day differentiation regimen. The presence of syncytia was confirmed by staining the cell membrane with wheat germ agglutinin (WGA) and Hoescht 33342 to visualize the nuclei (Figure 1E). ST-3D aggregates exhibited multinuclear structures with centrally clustered nuclei, whereas ST-2D cells displayed both multinuclear and mononuclear cells. TSCs continually proliferate in culture, as demonstrated by widespread Ki-67 positive staining, a macaque trophoblast marker of cell proliferation^40^, observed in nearly all TSCs (Figure 1F). The presence of Ki-67 positive mononuclear cells within ST-2D cultures further suggests that the ST-2D culture paradigm only partially commits cells to differentiate, whereas the ST-3D aggregates display fewer Ki-67 positive cells that are predominantly located at the periphery of the aggregate.

### TSCs and Differentiated Trophoblasts Express Trophoblast Genes

Classic trophoblast gene expression markers were evaluated for their presence in TSCs and pri-CTB by RT-PCR (n=4 cell lines). Both pri-CTB and TSCs expressed the hallmark epithelial trophoblast gene, KRT7, as well as transcription factors TEAD4, TFAP2C, and TP63 (Figure 2A). Variable CDX2 and ELF5 gene expression was observed in the TSC and pri-CTB lines, where CDX2 was not detected in cells from rh090419 and ELF5 was not detected in cy091318 TSCs. Expression of the macaque homolog of human HLA-G expression, macaque placental MHC class I molecule Mamu-AG^34^, was detected in all cells. To confirm RT-PCR results and to more comprehensively survey the transcriptome, RNA-seq was performed to profile gene expression of primary trophoblasts, reprogrammed TSCs and differentiated TSCs derived from the rh121118 line. RNA-seq revealed distinct gene expression profiles for each cell type, and unsupervised hierarchical clustering by expression level resulted in segregation of primary versus *in vitro* derived trophoblasts (Figure 2B). Gene expression values, represented as transcripts per million, of selected trophoblast-related genes displayed in Figure 2B are provided in Supplemental Table S2. TSCs more highly expressed trophoblast transcription factors including TP63, GATA3, GATA6 and TEAD4, whereas pri-CTB more highly expressed GCM1, CSH1 and 4, ID2, ELF5, GATA4 and a syncytium induction gene, ERVFRD1. The pri-ST cells that were differentiated *in vitro* displayed expression of classic ST markers including PLAC1, PLAC8, and Mamu-AG. By comparison, the ST-3D cells were enriched in genes representative of an early post-implantation placenta, as CGA and CGB expression was highly represented in comparison to pri-ST. ST-2D cells had expression profiles similar to both TSCs and ST-3Ds, with markedly high expression of TFAP2A, HAND1 and ESRRB, suggesting that these transcription factors may underlie initial commitment to ST differentiation.

**Figure 2.**
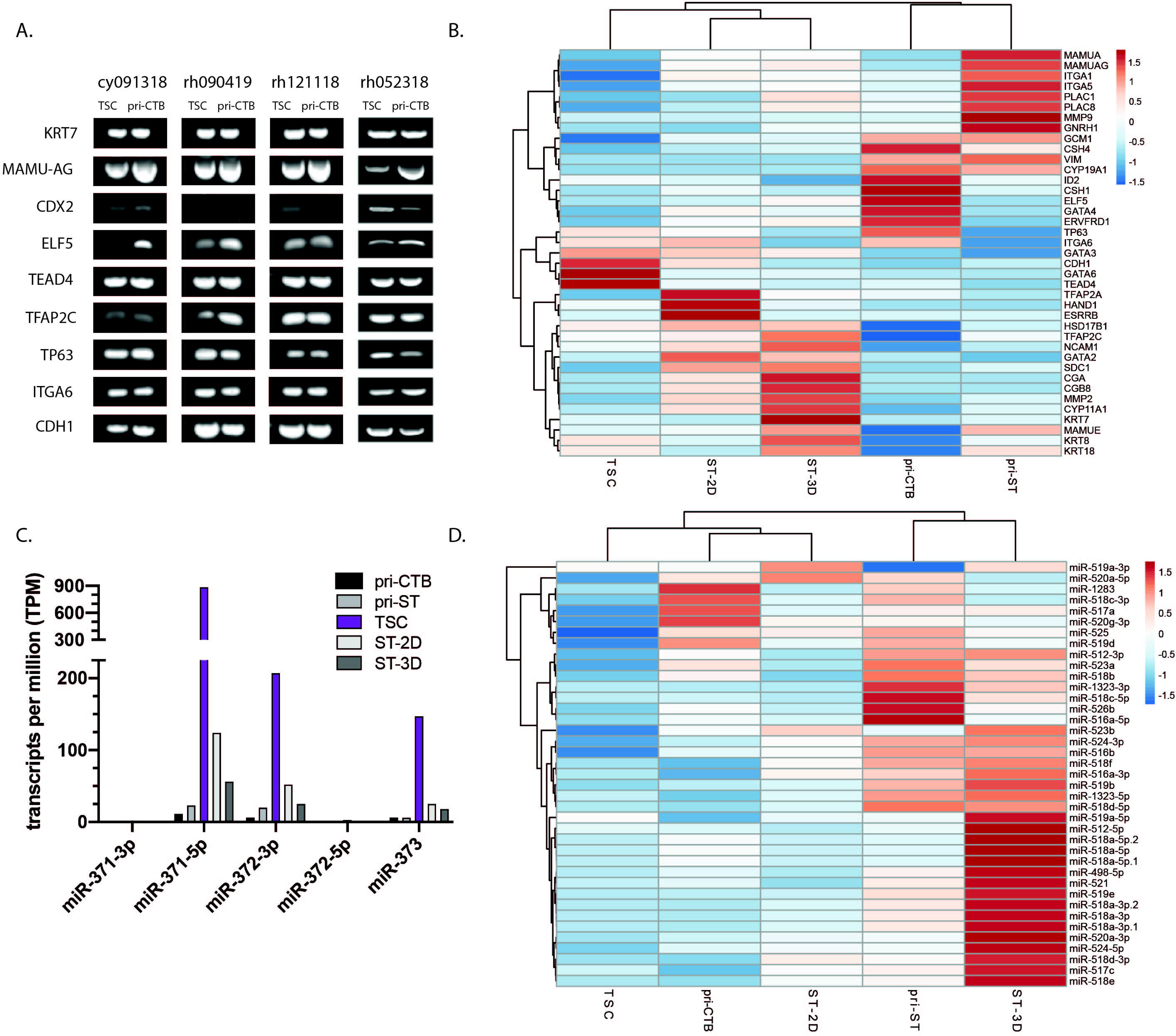
Expression of trophoblast marker mRNAs and miRNAs in parental primary villous cytotrophoblasts, reprogrammed TSC, and differentiated derivatives of TSC. A) RT-PCR products for trophoblast markers in four TSC lines versus isolated pri-CTB. B) RNA-seq profile of primary cells compared to derived rh121118 trophoblasts. Rows are mapped by correlation and columns by Euclidean distance. For panel B and D, the number of transcripts per million is scaled across rows with the red portion of the color scale bar representing higher values and blue showing lower values. C) miR-seq expression profile represented by transcripts per million (TPM) of miR-371-3 cluster miRNAs. D) miR-seq expression profile of chromosome 19 microRNA cluster (C19MC) miRNAs. Columns and rows are mapped as described for Panel B.

### Placental miRNA Cluster Expression

Primate trophoblasts uniquely express pregnancy-associated miRNA clusters^41, 42^, hence miR-seq analysis was performed to comprehensively survey expression of the miR-371-3, chromosome 19 microRNA cluster (C19MC), and chromosome 14 microRNA cluster (C14MC). The miR-371-3 is conserved in mammals and predominantly expressed in the placenta and embryonic stem cells with roles in cell cycle regulation^41, 43–47^. miR-seq analysis of the miR-371-3 cluster revealed elevated expression of this cluster’s miRNAs in TSCs compared to all other trophoblasts (Figure 2C, Supplemental Table S3). miR-371-5p expression was ∼800-times higher in TSCs compared to pri-CTB, with expression also detected in ST-2D and ST-3D cells. When comparing C19MC expression, the pri-CTB clustered with TSC and ST-2D expression levels while the pri-ST and ST-3D clustered together (Figure 2D). The ST cells, regardless of origin, highly expressed members of the C19MC (Figure 2D). Rhesus fibroblasts were also sequenced in parallel, and as expected the placenta-specific miRNAs were detected at low to undetectable levels (0 to <5 TPM; Supplemental Table S3). C14MC miRNAs were not detected in TSC or differentiated trophoblasts, whereas primary trophoblasts and fibroblasts weakly expressed C14MC miRNAs (Supplemental Table S3).

### TSCs Display Methylation Signatures Characteristic of Trophoblasts

Distinct methylation signatures at specific gene loci, such as ELF5, are a characteristic of trophoblasts^39^. Hemberger et al.^48^ have previously demonstrated that near the ELF transcription start site, the DNA of human trophoblasts is hypomethylated. ELF5 expression in mouse trophoblasts acts as a molecular switch between trophoblast proliferation and differentiation^28^. Methylation profiles were established for three TSC lines, three macaque blastocyst stage embryos, human BeWo choriocarcinoma cells and a rhesus fibroblast cell line within a 440 base pair amplicon containing an island of 11 CpG sites surrounding the macaque ELF5 start site (Figure 3A). Relative hypomethylation was observed in all 3 of the TSC lines analyzed, whereas embryos and BeWo cells displayed a hypomethylated profile, and, as expected, fibroblasts displayed a hypermethylated profile.

**Figure 3.**
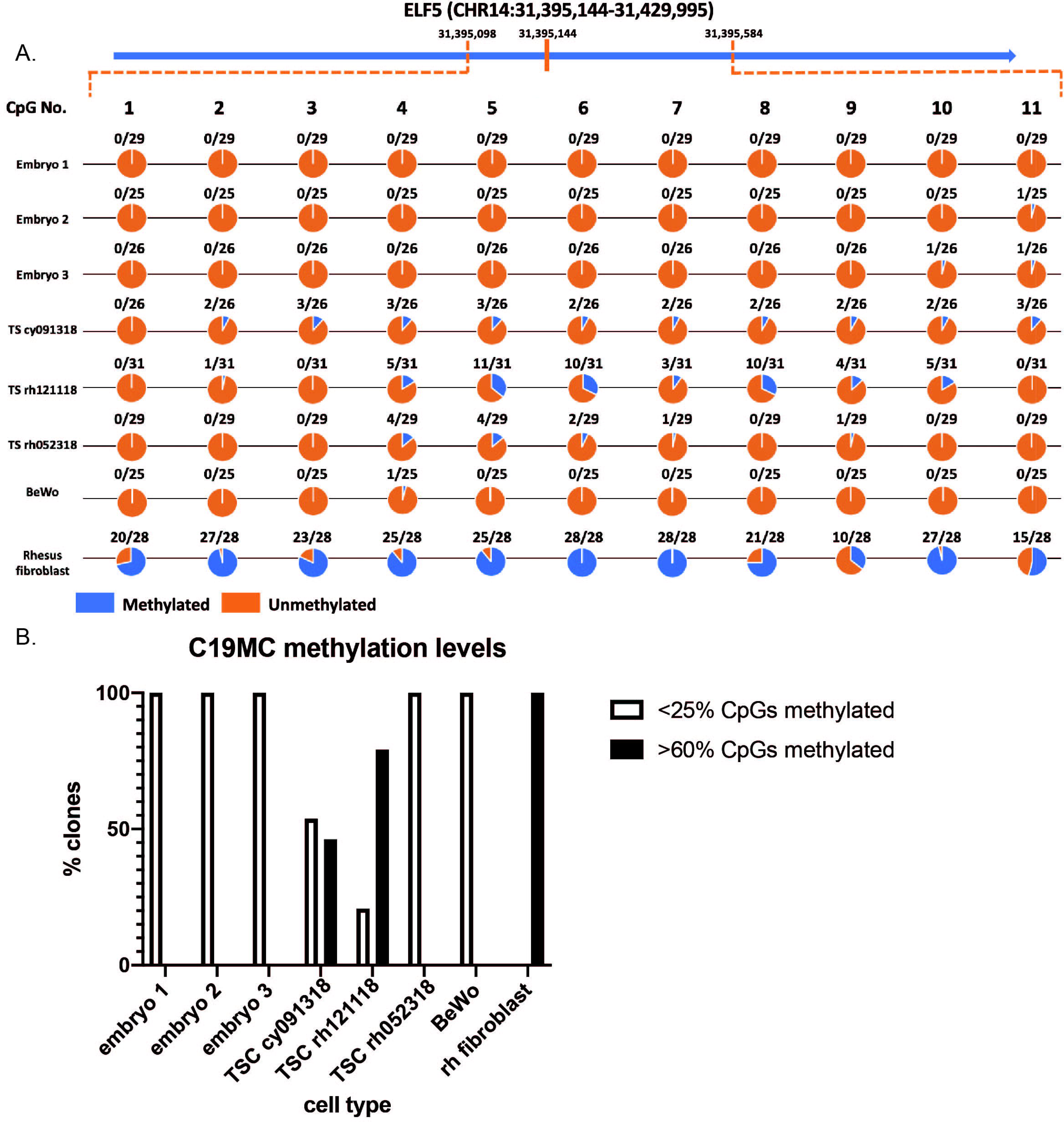
DNA methylation of ELF5 and C19MC gene regions. Methylation was evaluated in macaque blastocyst stage embryos (embryos 1-3), TSC lines, BeWo trophoblasts, and rhesus macaque fibroblasts. A) ELF5 methylation at 11 CpG sites in ∼18-30 clones for each cell type. Circles are colored to indicate the proportion of clones of clones either methylated (blue) or unmethylated (orange) at each CpG site. B) C19MC methylation at 34 CpG sites represented as the number of clones with either <25% or >60% methylation. All clones analyzed were either <25% or >60% methylated, representing an either hypo- or hyper-methylated allele, respectively.

Imprinted genes are vital to embryonic and placental development, and are monoallelically expressed from one parental allele while the other parental allele is methylated and not expressed^49^. The C19MC is positioned within an imprinted gene region where the methylation signature of the C19MC DMR1 in human placenta has been previously demonstrated by Noguer-Dance et al.^50^. Here, methylation of C19MC DMR1 was assessed similar to ELF5 to determine whether imprinting was maintained. Given that the C19MC DMR1 is maternally imprinted, it would be expected that 50% of the DNA clones would be hypermethylated and 50% of the clones would be hypomethylated representing the maternal and paternal allele, respectively, if imprinting was maintained. TSC lines varied in the degree of methylation: TS rh052318 cells were nearly all hypomethylated, while the TS cy091318 and TS rh121118 lines cells displayed both hypo- or hyper-methylated alleles (Figure 3B). In comparison, hypomethylation was observed in both macaque blastocyst stage embryos and BeWo cells, whereas rhesus fibroblasts which do not express C19MC miRNAs were predominantly hypermethylated. Furthermore, the lack of complete hypermethylation supports the expression of C19MC miRNAs observed in the TS rh121118 cells (Figure 2D).

### Trophoblasts and Differentiated ST Protein Expression

Trophoblast protein marker expression was validated by immunocytochemistry. Trophoblasts highly expressed KRT7/8, while vimentin was not detected (Figure 4A, see Supplemental Figure S2 for IgG control images). Interestingly, a classic trophoblast transcription factor, AP2, was present in both TSCs and ST cell types. Macaque ST differentiation markers, macaque chorionic gonadotropin (mCG) and Mamu-AG, displayed more intense staining in ST-3D aggregates compared to ST-2D cells (Figure 4A). Differentiated trophoblast marker expression thus further supports the induction of TSC differentiation to ST.

**Figure 4.**
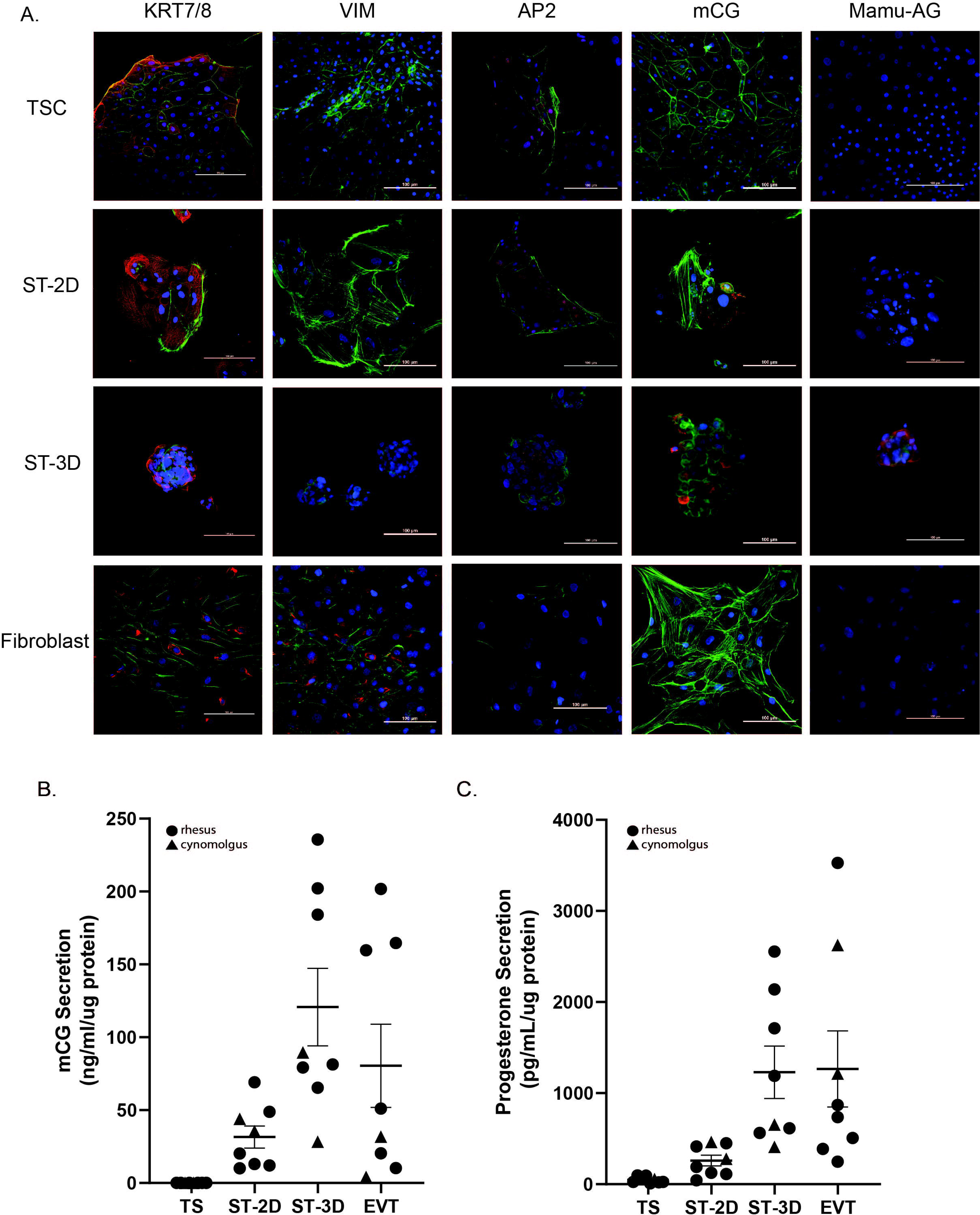
Differentiated trophoblast display protein markers of syncytium formation. A) Representative observations for immunocytochemistry against KRT7, Vimentin, AP2, mCG and Mamu-AG is represented in TSC, ST-2D, ST-3D and rhesus fibroblasts. For all panels, the specific marker of interest is shown in red, f-actin in green and the nuclei are stained with DAPI in blue. Scale bar denotes 100 µM. Secretion of mCG (B) and progesterone (C) by TS, ST2D, ST3D, and EVT cells was normalized to the amount of protein. Data is represented as the mean +/− the standard error of the mean (SEM). Data for rhesus macaque cells is indicated by circles, whereas data for cynomolgus macaque cells is indicated by a triangle. Significance was determined using a Kruskal-Wallis test with Dunn’s correction.

### ST cells secrete pregnancy hormones

The production of hormones, such as chorionic gonadotropin and progesterone, by trophoblasts is essential for the establishment and maintenance of pregnancy^51^. Secretion of mCG was significantly upregulated in ST-3D (120 ng/mL/µg protein) and EVT (80 ng/mL/µg protein) media, at ∼4000-fold higher levels in comparison to TSCs (0.02ng/ml/µg protein) (Figure 4B, p <0.007 and p <0.0001, respectively). ST-2D (31 ng/ml/µg protein) was ∼1500-fold higher than TSCs, but not significantly different (p=0.07). The levels of mCG detected in TSC conditioned media were near the lower limit of detection. Of note, the quantity of mCG secreted by EVTs varied quite substantially across cell lines. Progesterone secretion was also ∼1000-fold higher in ST-3D (1230 pg/mL/µg protein) and EVT (1265 pg/mL/µg protein) cultures compared to TSCs (48 pg/mL/µg protein; p<0.0003 and p <0.0006, respectively) and ∼5-fold higher in ST-2D (260 pg/mL/µg protein) media compared to TSC media (Figure 4C). Similar to mCG secretion, the levels of progesterone secretion by EVTs were similar to that of differentiated ST-3D cells.

### EVT Differentiation from TSCs

TSCs were cultured under the EVT differentiation paradigm as described by Okae et al.^29^, with the exception that the EVT culture medium either included or excluded the ALK-5 inhibitor, A83-01. Distinct EVT morphologies were observed as early as day 6 of the EVT differentiation program, where in the absence cells of A83-01 cell colonies were more sprawling, wider and flattened in comparison to cells with A83-01 supplemented to the media (Figure 5A, rh121118 cells presented as representative example). EVT morphology varied across lines, but was more consistent within an individual line (representative images shown in Supplemental Figure S4). To determine if the extracellular matrix influenced macaque differentiation capacity, TSCs were cultured in EVT medium with or without supplementation of A83-01 and were plated onto Matrigel. These cells, however, retained a more TSC like morphology (representative images in Supplemental Figure S4). Whereas the TSC and ST-2D cultures were supported on col IV-coated plastic wells or glass coverslips, EVT cell growth and adherence was not well-supported on col IV-coated glass. Thus, flow cytometry was used to determine trophoblast marker expression rather than immunocytochemistry.

**Figure 5.**
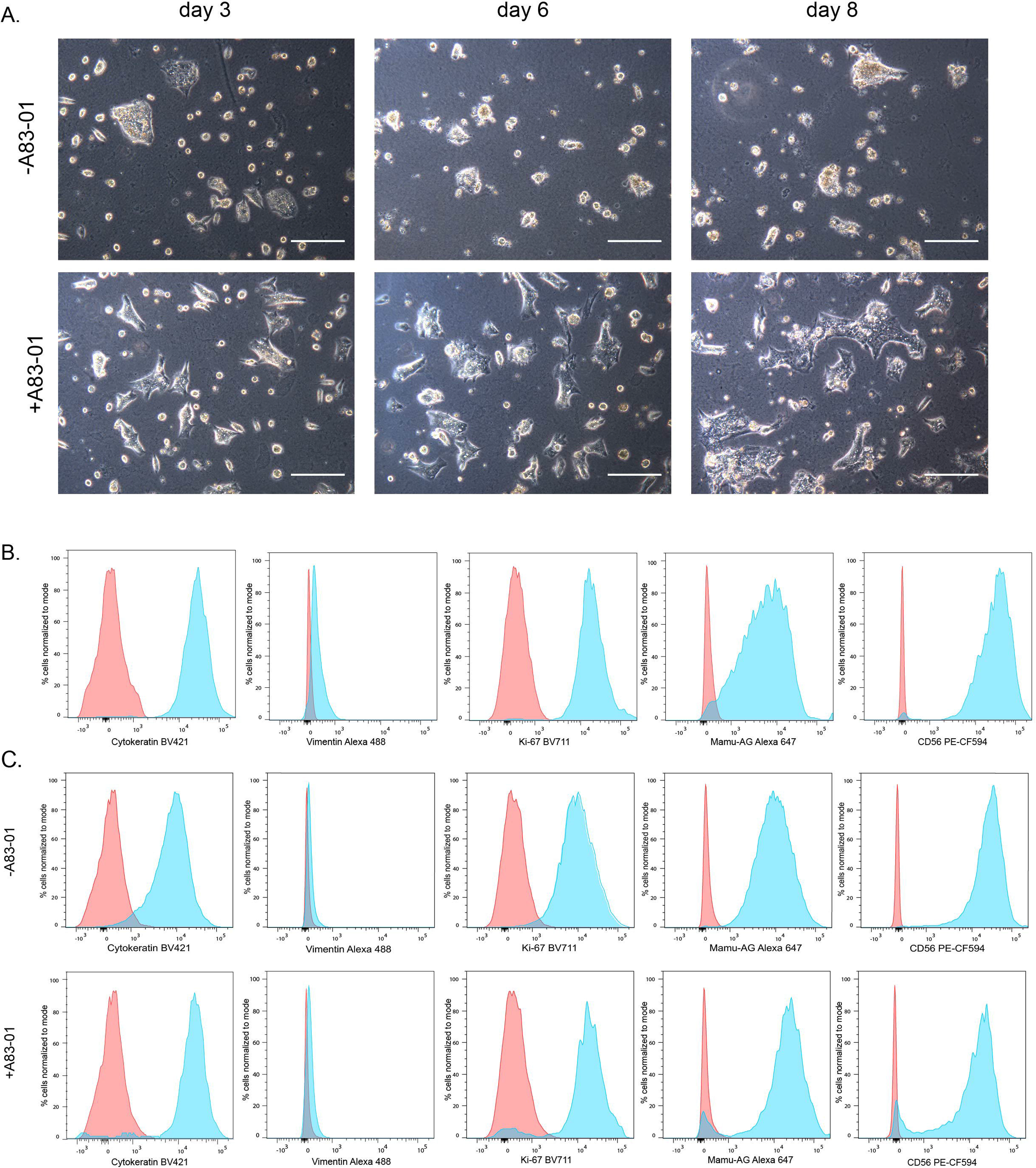
TSCs differentiate to an EVT-like cell type. A) rh121118 TSCs cultured in EVT medium with or without supplementation of A83-01 (denoted as +A 83-01 or -A83-01). Morphology representative of EVTs on day 3, 6 and 8 of culture. Scale bar denotes 250 µM. Flow cytometry of TSCs (B) and EVTs (C) against proteins cytokeratin, vimentin, Ki-67, Mamu-AG, and CD56. The proportion of cells that are stained (blue) or unstained (red) are represented as the % cells normalized to mode and the x-axis is the fluorescent intensity of the fluorophore.

Flow cytometry analysis was performed for all eight TSC lines, three EVT cultures each generated with or without media supplementation of A83-01, and a rhesus fibroblast cell line to serve as a negative control for trophoblast marker expression. TSCs were 70.8%-98.53% cytokeratin 7/8-positive, 97.0-99.9% vimentin-negative, 98.6-99.7% Ki-67-positive, 87.3-99.9% Mamu-AG-positive, 98.76-100% CD56-positive (Figure 5B, Supplemental Figure S4A). In comparison, rhesus fibroblast cells were 80.8% cytokeratin 7/8 positive, 97.9% vimentin positive, 93.84% Ki-67 positive, and 97.39% CD56 positive (Supplemental Figure S4B). Rhesus fibroblasts stained positive for Mamu-AG, however, with distinctly lower mean fluorescent intensity than that of TSCs or EVTs (Supplemental Figure S4B). EVT marker expression did not differ when cultured with or without inclusion of A83-01 in the culture media with at least 94% positive-cytokeratin, Ki-67, Mamu-AG and CD56 cells and 99% vimentin-negative cells (Figure 5, Supplemental Figure S5). In one experiment, to assess the support of differing extracellular matrices, TSCs of rh121118 were plated on Matrigel coated wells and cultured in EVT medium with or without A83-01 supplementation to compare to those cultured on col IV. EVT morphologies varied between cultured conditions (EVTs on col IV are shown in Figure 5A and EVTs on Matrigel are shown in Supplemental Figure S3B), however, flow cytometry revealed similar populations of Mamu-AG and CD56 positive cells regardless of the extracellular matrix (Figure 5C, Supplemental Figure S5B).

### Single Cell RNA-sequencing of early and established TSC passages

Single-cell RNA-sequencing (scRNA-seq) was performed to assess the heterogeneity of TSC cultures and to evaluate changes in gene expression from an early passage (p2) or an established TSC passage (p10). A total of 11,311 cells (p2: 6,052 cells, p10: 5,259 cells) were sequenced, in which 73,091 mean reads were detected per cell and the median number of genes per cell was 1,468. The Louvain Modularity Optimization (LMO) algorithm^52^was used to cluster cells on the number of nearest neighbors (based on cellular gene expression levels) and the clusters were plotted using a dimensionality reduction method, t-SNE. The gene expression of p2 cells (Figure 6A) clustered more tightly, indicative of a more homogeneous population than that of p10 cells (Figure 6B). There were no significantly differentially expressed genes with a fold-change >2 within the nine p2 clusters, whereas p10 cell gene expression segregated into 11 clusters, of which seven clusters had at least one significantly differentially expressed gene indicated in the legend identifying each cluster (Figure 6 AB; Supplemental Table S4).

**Figure 6.**
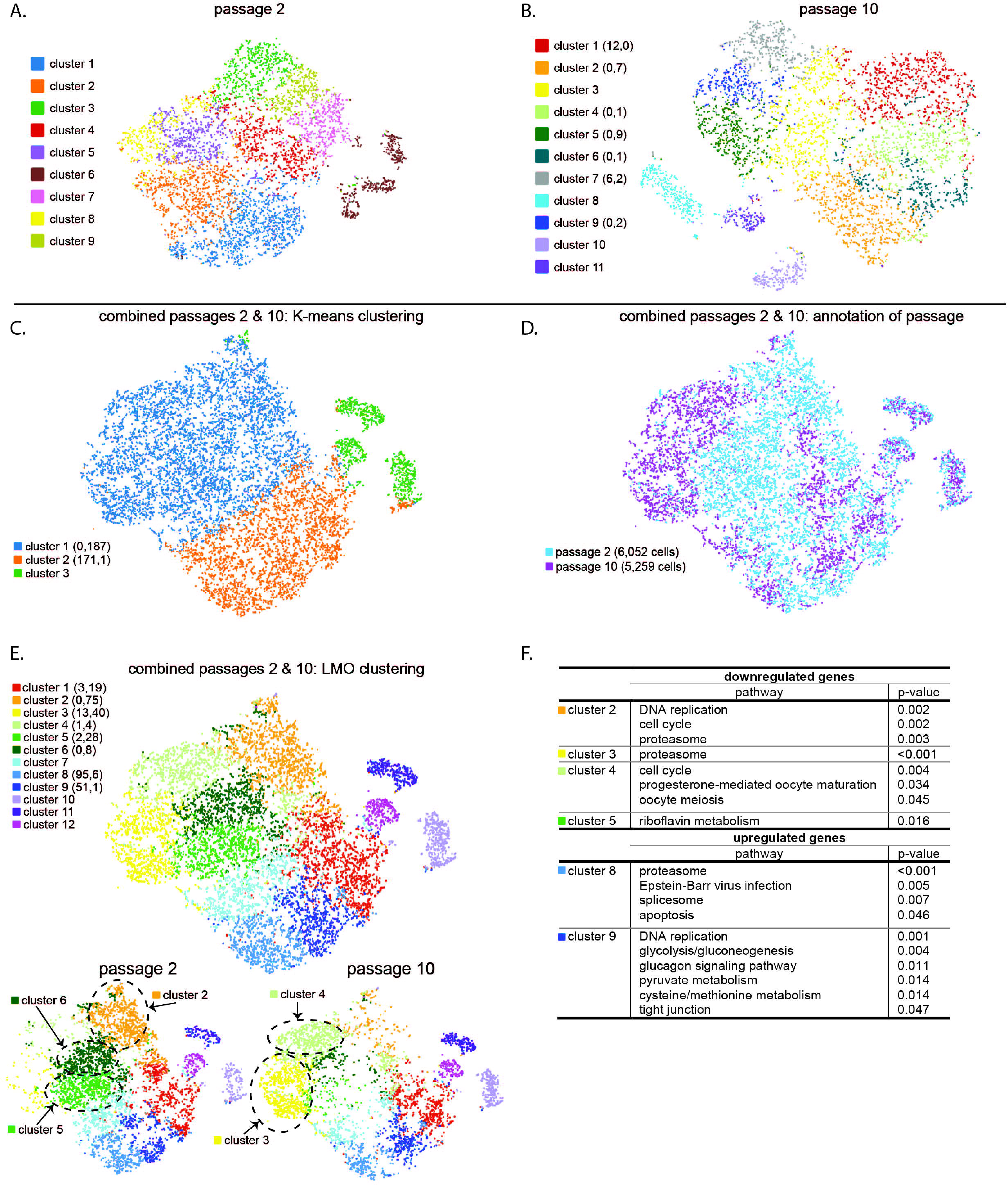
TSC gene expression evaluated by scRNA-Seq. t-SNE distribution of cellular gene expression at passage 2 (p2, A) and passage 10 (p10, B). The number of significantly up or down regulated genes in this and other panels appear in parentheses next to the cluster number. C) Gene expression by K-means clustering analysis of combined p2 and p10 cells. D) Identification of the distribution of individual p2 and p10 cells within the t-SNE distribution with the number of individual cells sequenced for each passaged denoted in parentheses. E) Gene expression of combined p2 and p10 cells analyzed by Louvain Modularity Optimization algorithm and plotted using t-SNE segregated cells into 12 clusters. Individual cells of p2 and p10 of the combined analysis are represented below. Clusters where cells were more highly represented from the early or more established TSC passage are indicated by gray circles. F) Human KEGG analysis of significantly altered pathways by up or down regulated genes within specific clusters from panel E. Up and down regulated genes were analyzed using ENRICHR and those pathways with significant adjusted p-values are represented.

A K-means analysis, which is a more conservative method for clustering, was performed on combined TSCs of p2 and p10. The K-means analysis revealed three distinct cell clusters (Figure 6C), where an inverse differential expression profile was observed between clusters 1 and 2 with similar genes either up or down regulated between the clusters. Cluster 1 had 187 downregulated genes, whereas cluster 2 had 171 upregulated genes and 1 downregulated gene (Figure 6C, Supplemental Table S5). Highly differentially expressed genes in between K-means clusters 1 and 2 include NUCB2, UBB, TECR, PMEL, UBC, APOE, MRPS31, PRDX2, and LDHA (Figure 7A), where each of these genes were downregulated in cluster 1 cells and significantly upregulated in cluster 2 cells. The individual expression profiles for the top differentially expressed genes identified by K-means are illustrated in Figure 7A.

**Figure 7.** scRNA-seq expression of individual genes within the t-SNE distribution early and established TSC passages. Top differentially expressed genes in common between clusters identified by K-means clustering analysis (A) or LMO algorithm (B). C) Representation of individual trophoblast markers. Scale bar represents the range in Log2 expression for the marker for all panels.

Clustering analysis using the LMO algorithm segregated cells of both passages with similar expression profiles into 12 clusters that are represented by a t-SNE distribution plot with individual cells denoted by passage (Figure 6D) or by cluster (Figure 6E). Differentially expressed genes of each cluster are listed in Supplemental Table S6. Cells of different passages clustered among each other or were segregated, as the cells that comprised clusters 3 and 4 were predominantly cells of p10 and cells of clusters 2, 5, and 6 were largely of p2 (Figure 6E). Using the Human Kyoto Encyclopedia of Genes and Genomes (KEGG) database, pathway analysis of cluster 2 downregulated genes revealed that the pathways significantly altered included DNA replication, cell cycle and proteasome pathways (Figure 6F). Gene ontology (GO) of biological processes for downregulated genes of cluster 2, 4, and 6 featured mitotic cell cycle transition as a common term between them, whereas, Wnt signaling pathway was a significantly altered process of downregulated genes of clusters 2, 3, 4 and 5. The alteration of similar pathways by multiple clusters is likely a reflection of a subset of similarly differentially expressed genes that are shared across clusters. For example, clusters 1, 2 and 4 significantly downregulated TOP2A, CDK1, GTSE1 and UBE2C, whereas cluster 5 cells had upregulated expression TOP2A and GTSE1. Of note, these genes represent a few of the differentially expressed genes by which the clusters segregate. The distributions and expression levels of selected genes are shown in Figure 7B. Interestingly, members of the bone morphogenetic protein (BMP) family have been shown to drive human ESC differentiation to trophectoderm^18^, and here TSCs of cluster 3 had upregulated expression of BMP7 (Figure 7B). When evaluating expression of classic trophoblast markers in individual cells, the expression of these markers was not significantly different between clusters and rather were represented throughout the t-SNE distribution (Figure 7C).

## Discussion

The present study demonstrates the derivation of macaque TSCs that not only express key trophoblast features, but also are capable of continued proliferation as well as differentiation to ST was achieved. Using the paradigm established for the derivation of human TSCs, we have shown that macaque TSCs can be similarly derived from pri-CTB that are capable of tightly controlled proliferation and differentiation. This is in contrast to previously reported generation of macaque TSCs from rhesus blastocysts^38^. Criteria for defining trophoblast cell types as described by Lee et al.^39^ include the expression of trophoblast protein markers, placental MHC class I expression, hypomethylation of ELF5, and expression of placenta-specific C19MC miRNAs. Expression of each of these criteria is demonstrated for macaque TSCs and differentiated ST. Of note, HLA-G is not expressed in the macaque placenta but rather Mamu-AG serves as the MHC class I homolog^33^ and is expressed as expected in ST and EVT cells. Moreover, macaque ST-3D and EVT cells secreted high levels of mCG and progesterone, similar to human primary cultures of ST and EVT^53^. Altogether, trophoblast marker expression specific to TSC, ST, and EVT in the macaque TSC model was similar to that described in human TSC models^28–30^.

Under standard culture conditions, macaque pri-CTB will spontaneously differentiate to multinuclear syncytia^54, 55^, similar to human trophoblasts^20^. With the TSC “reprogramming” paradigm, pri-CTB switch to a mononuclear, proliferative state within a few days of culture in TSC medium that promotes Wnt activation and EGF signaling, while inhibiting TGF-ß signaling. As early as 24 hours after plating, primary trophoblasts cultured in basal medium readily form syncytia and do not continue to proliferate. Therefore, a molecular switch likely occurs at this time to support cell renewal. Differentially expressed genes observed in the scRNA-seq data are predominantly genes that regulate the progression of the cell-cycle and ultimately cellular proliferation. Genes such as topoisomerase 2 alpha (TOP2A) and cyclin dependent kinase-1 (CDK1) are down regulated in cluster 1, 2 and 4 cells (both p2 and p10 cells), whereas TOP2A expression was significantly increased in cluster 5 cells (predominantly p2 cells). TOP2A aids in the relaxation of DNA supercoils during replication, and is expressed in a cell-cycle dependent manner with increased levels in mid-S phase through mitosis^56^. Likewise, CDK1 is essential for transition through the cell cycle^57^. The differential expression of these genes and others with roles in cell replication suggest that the scRNA-seq clusters reflect cell populations that are either actively proliferating or that are more quiescent. Interestingly, BMP7 was highly expressed in cluster 3 which was predominantly comprised of established TSCs. Unlike human embryonic ESCs, macaque ESCs do not respond to BMP4 treatment to promote differentiation to trophectoderm^58, 59^, however, it is plausible that this cluster represents established TSCs that are positioned for differentiation. A more conservative analysis, K-means analysis, revealed a distinct axis of up or down-regulated genes between clusters 1 and 2, including NUCB2, APOE, PRDX2, and LDHA, that have previously been shown to have roles in trophoblast function^60–64^. The roles of these genes in TSC function and maintenance, however, largely remains unclear. Importantly, irrespective of expression analysis method, no trophoblast markers were differentially expressed across the clusters. For example, cells that expressed markers such as TFAP2C or KRT18 were distributed throughout the t-SNE plot.

Macaque TSCs display gene and protein expression as well as DNA methylation features similar to human TSCs. Based on flow cytometry and immunocytochemistry findings, all TSC lines were predominantly cytokeratin-positive and vimentin-negative. While KRT7 expression is a hallmark of human trophoblasts^39^, macaque TSCs displayed higher RNA expression of KRT18 compared to KRT7. Macaque TSCs also expressed trophoblast transcription factor genes TEAD4, TP63 and TFAP2C similarly as reported for human TSCs^28, 29^. Weak expression of CDX2 was observed in pri-CTB and TSCs by RT-PCR, and comparison to RNA-seq data was not feasible given that the CDX2 transcript is not annotated in the rhesus macaque reference genome used for sequencing alignment (BCM Mmul_8.0.1/rheMac8). Similarly, ELF5 displayed lower expression in TSCs compared to pri-CTB by RNA-seq, with a limited detection in TSCs by scRNA-seq analysis. Due to the limited characterization of expression of macaque trophoblast transcription factors, histological and quantitative gene expression analysis of CDX2 and ELF5 in macaque TSCs and placentas is needed to determine if these markers are conserved between humans and macaques. At the genomic level a hypomethylated pattern was observed near the ELF5 gene start site, similar to observations in human TSCs^28–30^ and placentas^48^. Moreover, this is the first study to evaluate C19MC methylation in macaque trophoblasts and embryos. As the C19MC is a maternally imprinted gene region, it is expected that the maternal allele is hypermethylated while the paternal allele is hypomethylated^50^. Although the maternal and paternal alleles were not determined in this study, it could be anticipated that a 1:1 ratio of hyper-to hypo-methylated DNA clones from TSCs would be observed. Methylation at the C19MC DMR1 locus, however, varied across TSC lines where the rh052318 TSCS were largely hypomethylated compared to TSCs of cy091318 which had near a 50:50 ratio of hypo- to hyper-methylated DNA clones. Future analysis of additional TSC lines and clones is needed to determine the maintenance of C19MC imprinting in the derivation and maintenance during cell culture, with subsequent analysis of the relationship between C19MC methylation and miRNA expression levels.

An important hallmark of human and NHP trophoblasts is the expression of placental miRNA clusters^35, 41, 42^. TSCs exhibited an upregulation of the miR-371 cluster miRNAs. The miR-371 cluster is predominantly expressed in the mammalian placenta^45^ and is highly expressed in ESCs^47, 65^ with decreased expression upon ESC differentiation^47^. In a previous study, we have shown that miR-371 cluster expression tends to be higher in first trimester macaque placentas compared to term placentas^35^. Knock-out of the orthologous mouse miRNA cluster, the miR-290 cluster, suggested that the miRNAs of this cluster have a role in the cell cycle and in maintaining mitotic divisions of trophoblast progenitor cells^66^. Zhou et al.^67^ reported a dynamic relationship between the interaction of the miR-371 cluster and the Wnt/ß-catenin signaling pathway, a pathway implicated in stem cell maintenance and cancer cell proliferation. Upon inhibition of glycogen synthase kinase-3 (GSK-3) and suppression of ß-catenin ubiquitination, to stimulate Wnt/ß-catenin signaling, miR-371 cluster expression was increased through induction by LEF1/ß-catenin following activation of Wnt signaling. Moreover, miR-372 and miR-373 induced ß-catenin expression as well as increased expression of cell cycle regulators c-Myc, c-Jun, and cyclin-D1^67^. Hence, the presence of the GSK-3 inhibitor, CHIR99021, in primate TSC culture media to activate Wnt signaling also promotes the placental miR-371 cluster expression, which in turn acts as a positive-feedback mechanism to promote Wnt signaling. The interaction between Wnt signaling and miR-371 is likely a key factor to the stemness and continued proliferation of primate TSCs.

Upon culture of TSCs with forskolin, ST differentiation occurs with high production and secretion of mCG. The ST-3D culture was effective at inducing syncytium formation and expression of ST markers. For instance, ST-3D aggregates demonstrated higher RNA expression for CGA and CGB and trended towards greater secretion of mCG in culture media compared to ST-2D cultures. Moreover, ST-2D cultures retained a subset of mononuclear TSCs and expressed a gene profile that may reflect initial or partial commitment to ST differentiation. More importantly, the observance of mCG secretion is indicative of a primitive or early syncytium. Macaque CG secretion is transient, with serum and urine concentrations peaking between day 21-25 post menses or days 8-12 post-conception^37, 68, 69^. Given that the pri-CTB were isolated from gestation day 40-74 placentas, there was limited to no expression of CGA and CGB genes in pri-CTB or pri-STB. The secretion of mCG by differentiated ST-2D and ST-3D demonstrates that the TSCs were reprogrammed to a more primitive or early gestation trophoblast population. Thus, the macaque TSCs may be suitable for modeling events of early placental development.

The differentiation conditions to derive macaque EVTs from TSCs requires further optimization with subsequent development of criteria to define macaque EVT cells. Unlike human HLA-G, the MHC class I homolog, Mamu-AG, is not exclusive to macaque EVTs but rather is expressed in both ST and EVT cell populations^33^. The cell adhesion protein, CD56 or NCAM1, is expressed by macaque trophoblasts within the cytotrophoblastic shell and endovascular EVTs^70^. Macaque TSC-derived EVTs in the present study expressed both Mamu-AG and CD56, however, different morphologies were observed in the presence or absence of the ALK5-inhibitor A83-01. The ALK5 receptor binds TGF-ß and activin, hence the small molecule inhibitor A83-01 acts to inhibit this binding, which prevents downstream activation of SMAD signaling pathways, and altogether has been shown to inhibit the epithelial-mesenchymal transition (EMT) induced by TGF-ß signaling^71^. An EMT is thought to occur during EVT differentiation as EVTs transition from an epithelial to mesenchymal cell type with loss of their attachment to a basement membrane, allowing for migration and invasion into the maternal decidua^72, 73^. DaSilva-Arnold et al.^73^ observed increased gene expression of TGFB1 and TGFB2 in human EVTs compared to primary villous cytotrophoblasts suggesting that TGF-ß expression is intrinsic to human EVTs. Supplementation of exogenous TGF-ß1 to human primary cultures reduced proliferation and promoted formation of multinuclear cells^74^. In the context of TSC culture, TGF-ß inhibition by A83-01 may be required to promote proliferation, while it may inhibit the EMT necessary for EVT formation. The inhibitor was not included in EVT differentiation media in the human TSC studies described by Haider et al.^28^ and Turco et al.^30^, but was included in the EVT differentiation paradigm described by Okae et al.^29^. TGF-ß is secreted by decidual NK cells^75^, although the concentration of the cytokine that EVTs are exposed to depending on their migration into the decidua may contribute to differing cellular phenotypes. Future studies should focus on evaluating the concentration of A83-01 supplementation to determine whether there is a dose-dependent effect on EVT marker expression. Moreover, it is plausible that macaque EVTs generated with or without inhibition of TGF-ß represent EVTs at different stages of differentiation and further characterization of markers for macaque progenitor, interstitial and endovascular EVTs is needed to define *in vitro* derived EVTs.

In conclusion, macaque pri-CTB can be driven to TSCs utilizing derivation conditions previously described by Okae et al.^29^ for human embryonic trophectoderm and placental pri-CTB. The capacity of macaque TSCs to self-renew and differentiate to ST under a tightly-controlled 3-D culture system offers a platform for modeling primate trophoblast differentiation and development in vitro. More importantly, in the macaque model experimental perturbations can be first evaluated within the in vitro TSC model with subsequent translation to macaque embryos and *in vivo* pregnancy, allowing for comprehensive study at the cellular and organismal level. For instance, the macaque TSC model can be directly used to perturb gene functions underlying trophoblast differentiation processes, evaluate trophoblast response to experimental infection, or assess the efficacy of placental therapeutics tailored to specific trophoblast cell types. The NHP TSC model could be pivotal to understanding mechanisms of early placental development leading to the development of treatments or therapeutics to improve human pregnancy health.

## Methods

### Subjects

Pregnant rhesus and cynomolgus macaques were obtained from the colony maintained at the Wisconsin National Primate Research Center (WNPRC). This study was performed in strict accordance with the recommendations by the National Research Council Guide for the Care and Use of Laboratory Animals in an AAALAC accredited facility (WNPRC). Experimental procedures were approved by the University of Wisconsin-Madison Institutional Animal Care and Use Committee (protocols: g005061, g005691). Macaque placentas were collected between gestation day 40-74.

### Placenta Cell Dissociation

First trimester placental tissue was dissociated to obtain primary villous cytotrophoblasts (pri-CTB) utilizing a trypsin/DNase and Percoll gradient isolation method as previously described^54^. Following gradient isolation, the cell pellet was then resuspended in DMEM. A fraction of pri-CTB for the rh121118 line were differentiated into primary ST (pri-ST) by plating ∼1.5-2×10^6^ cells per 60 mm dish and grown in DMEM for 72 h. The pri-ST and pri-CTB were collected into TRIzol Reagent (Invitrogen, ThermoFisher Scientific) for RNA isolation.

### Generation of Trophoblast Stem Cells

Macaque TSCs and differentiated trophoblast cells were derived utilizing similar methods as those described by Okae et al.^29^ to generate human TSCs and subsequently differentiated trophoblast cell populations. Primary-CTBs were plated at a density of 2-5×10^5^ cells per well of a 6-well plate coated with 5 µg collagen IV (col IV, Corning, cat no: 354233) and maintained in trophoblast stem cell (TSC)-medium described by Okae et al.^29^ consisting of DMEM/F12 supplemented with 0.1 mM 2-mercaptoethanol (Gibco, ThermoFisher Scientfic, cat no: 31350010), 0.2% FBS (Peak Serum, cat no: PS-FB1), 0.5% Penicillin-Streptomycin, 0.3% BSA (Sigma-Aldrich, cat no: A19335G), 1% ITS-X supplement (ThermoFisher Scientific, cat no: 51500056), 1.5 µg/mL L-ascorbic acid (Wako Chemical USA, cat no: 013-12061), 50 ng/mL EGF (ThermoFisher Scientific PHG0313), 2 µM CHIR99021 (Tocris, cat no: 4423), 0.5 µM A83-01 (Tocris, cat no: 2939), 1 µM SB431542, 0.8 mM VPA (Sigma-Aldrich, cat no: P4535) and 5 µM Y27632 (Tocris, cat no: 1254). Cell cultures were maintained at 37°C and 5% CO_2_ and medium was replaced every 1-2 days. Once cells reached 80-90% confluency, cells were passaged by incubating with TrypLE Select (Gibco, ThermoFisher Scientific, cat no: 12604021) at 37°C for 15 min to lift the cells. Cells were plated at a density of 1-2.5×10^5^ cells per well of a 6-well plate coated with 5 µg col IV or 0.5-1×10^6^ cells per T75 flask coated with 25 µg col IV.

### In vitro Trophoblast Differentiation

TSCs were differentiated to ST utilizing either two-dimensional (ST-2D) or a three-dimensional (ST-3D) differentiation protocols as described by Okae et al.^29^. To obtain ST-2D cells, TSCs were plated at a density of ∼2×10^5^ cells per well of a 6-well plate coated with 1 µg collagen IV and maintained in ST-2D medium consisting of DMEM/F12 supplemented with 0.1 mM 2-mercaptoethanol, 0.5% Penicillin-Streptomycin, 0.3% BSA, 1% ITS-X supplement, 2.5 µM Y27632, 2 µM forskolin (Cayman Chemical, cat no: 66575-29-9) and 4% Knockout Serum Replacement (KOSR; ThermoFisher Scientific, cat no: 10828028). Medium was replaced on day 3 of differentiation. On day 6, cellular morphology was assessed and cells were collected. To obtain ST-3D cells, 2.5×10^5^ TSCs were seeded into a low binding T25 flask (Nunc, ThermoFisher Scientific, cat no: 169900). Suspended cells were maintained in 3 mL of ST-3D medium containing DMEM/F12 supplemented with 0.1 mM 2-mercaptoethanol, 0.5% Penicillin-Streptomycin, 0.3% BSA, 1% ITS-X supplement, 2.5 µM Y27632, 50 ng/mL EGF, 2 µM forskolin and 4% KOSR. An additional 3 mL of ST-3D medium was added on day 3 of differentiation. On day 6 of differentiation, the cell suspension was passed through a 40 µM filter to remove dead cells and debris. ST-3D aggregates were washed from the filter with DMEM/F12 medium and collected.

To differentiate TSCs to EVTs, TSCs were plated at a density of ∼1×10^5^ cells per well of a 6-well plate coated with 5 µg collagen IV per well. After adding the cell suspension to the well, Matrigel (Corning, cat no: 354234) was added at a final concentration of 2%. From day 0-3, cells were cultured in 2 mL of EVT medium comprised of DMEM/F12 supplemented with 0.1 mM 2-mercaptoethanol, 0.5% Penicillin-Streptomycin, 0.3% BSA, 1% ITS-X supplement, 100 ng/ml NRG1 (Cell Signaling, cat no: 5218SC) 2.5 µM Y27632, 4% KOSR and with or without the addition of 7.5 µM A83-01. On day 3 of culture, medium was exchanged with EVT medium lacking NRG1 with or without the supplementation of A83-01, and Matrigel was supplemented at a final concentration of 0.5%. On day 6 of culture, medium was replaced with 2 mL of EVT medium lacking NRG1 and KOSR, with or without the addition of A83-01, and Matrigel was supplemented at a final concentration of 0.5%. On day 8 of culture, cells were collected for analysis. EVT media were collected from cells that were cultured in the presence of A83-01 for hormone analysis.

### Cytogenetics

Karyotyping was performed by Cell Line Genetics (Madison, WI) using methods previously described^76^. In brief, cells were treated with 0.1 μg/ml colcemid for 40 minutes and then enzymatically treated to produce a single cell suspension. Cells were then incubated in 0.075M KCl, followed by fixation using Carnoy’s fixative (3:1 methanol:glacial acetic acid). Fixed cells were dropped and dried onto slides using a Percival environmental chamber and baked at 90 °C for 60 min. Chromosomes were then banded by utilizing a trypsin treatment followed by staining with Geimsa. Images were taken of metaphase cells using a GSL scanner (Leica) and analyzed with the aid of CytoVision software (Leica).

### RNA Isolation

Cells collected for RNA isolation were resuspended with 1 mL of TRIzol reagent (Invitrogen, cat no: 15596018, ThermoFisher Scientific) and frozen until RNA extraction. The TRIzol-cell mixture was layered onto a Phasemaker Tube (Invitrogen, ThermoFisher Scientific, cat no: A33248) and incubated at room temperature for 3 min followed by addition of 200 µL chloroform (MP Biomedicals, ThermoFisher Scientific, catalog no: ICN19400280) and spun at 16000 g at 4 °C for 15 min to allow separation. The aqueous layer was recovered, 500 µl 70% ethanol was added, and then transferred a RNeasy Mini Kit (Qiagen, ThermoFisher Scientific, cat no: 74104,) spin column and proper buffer washing steps were carried out following manufacturer recommendations including a 15 min on-column DNAse (Qiagen, ThermoFisher Scientific, cat no: 79254) treatment. RNA concentration was then quantified via Qubit Fluorometer using the RNA BR Assay Kit (Qiagen, ThermoFisher Scientific, cat no: Q10210,) and quality was evaluated using an Agilent 2100 BioAnalyzer (Agilent, cat no: G2939BA).

### cDNA Synthesis and RT-PCR

cDNA was synthesized using a SuperScript III First-Strand Synthesis System (Invitrogen, ThermoFisher Scientific, cat no: 18080-051). RNA from four cell preparations of each pri-CTB and corresponding TSC were combined with kit reagents and reactions were performed following manufacturer recommendations. PCR was performed by combining cDNA, GoTaq Hot Start Colorless Master Mix (Promega, cat no: M513B), RNAse-free water, and gene-specific primers (Supplementary Table S7). Negative control no reverse transcriptase cDNA reactions and no cDNA template were included for each template and primer pair. PCR reactions were performed using a BioRAD C1000 Touch instrument. The PCR products were then run on a 2% agarose gel and the bands were analyzed to determine gene expression.

### Next-generation RNA sequencing

Total RNA was isolated from each cell line derived from rh121118 for transcriptome and miRNA expression analysis. For transcriptome analysis, ∼1 µg total RNA was used as input for each cell type (pri-CTB, TSC, pri-ST, ST-2D, ST-3D, EVT and fibroblast) to generate cDNA libraries using a TruSeq Stranded mRNA library preparation kid (Illumina). Single-end 100 bp reads (1×100) were sequenced on an Illumina HiSeq 2500 instrument using a rapid run mode. Reads were trimmed using Skewer^77^, and then aligned to the rhesus macaque genome (BCM Mmul_8.0.1/rheMac8) using Spliced Transcripts Alignment to a Reference (STAR) software^78^, a splice junction aware aligner. Expression estimation was performed using RSEM (RNA-Seq by Expectation Maximization)^79^ and reported as transcripts per million (TPM). To survey miRNA expression, ∼100 ng of total RNA from each cell type was used as input to generate miRNA sequencing libraries using a QIAseq miRNA library kit (Qiagen). Single-end 100 bp reads (1×100) were sequenced on an Illumina HiSeq 2500 using a rapid run mode. To estimate known miRNA abundance, the miRNA-Seq workflow in miARma-Seq was used^80^. Accessions for known miRNAs were obtained from miRbase v22 (October 2018). Library construction and sequencing steps were performed by the University of Wisconsin Biotechnology Center.

### Immunocytochemistry

For TSC and ST-2D cultures, cells were cultured on col IV coated glass coverslips within a 6-well plate. TSCs were grown for 3-4 days prior to fixation, whereas ST-2D were fixed on day 6 of the differentiation paradigm. ST-3D aggregates were grown in suspension and on day 6 were plated on to col IV coated coverslips for 1-2 h to allow for attachment, immobilizing the aggregates for ease of handling and imaging. At the appropriate time points the cells were rinsed with PBS and fixed with 2% PFA for 10 min at room temperature. The coverslips with adhered cells were then washed twice with PBS and stored in PBS at 4 °C until processed. The samples were rinsed with PBS, permeabilized with 0.1% Triton-X 100 in PBS for 5 min, blocked using Background Punisher (Biocare Medical, cat no: BP974) for 10 mins, rinsed and exposed to primary antibodies for 1 h at room temperature by floating on a 75 µl drop, see Supplemental Table S8 for antibody and fluorescent conjugate information. The coverslips were then washed with PBS three times for 5 min each, and floated on drops containing the appropriate conjugated secondary antibodies and phalloidin conjugate for 45 min at room temperature followed by three washes for 10 min each at room temperature with TBST (Tris buffered saline containing 0.1% Tween-20). The coverslips were then treated with DAPI (Invitrogen, cat no: D3571; 1:10000 dilution) for 5 min at room temperature, rinsed with water, and mounted onto slides using Prolong Diamond Antifade Mountant (Invitrogen, cat no: 36970, ThermoFisher Scientific). Cells were imaged on either a Nikon A1R confocal or a Nikon Eclipse Epifluorescent microscope.

To visualize nuclei and cell membranes, cells were stained with wheat germ agglutinin (WGA) which binds to the carbohydrates on the cell surface. Cells on coverslips were generated as described above, and fixed samples were rinsed with PBS but not permeabilized. The samples were then stained with WGA conjugated with Texas Red (5 µg/ml) and Hoerscht 33342 for 10 min at room temperature, rinsed three times in PBS for 10 min each, followed by one rinse in water to removed salts prior to mounting with ProLong Diamond Antifade Mountant (Invitrogen, ThermoFisher Scientific, cat no: p36961) onto a glass slide. Cells were imaged on a Nikon A1R confocal microscope.

### Hormone Secretion Assays

Cell conditioned media were collected from confluent TSCs or at the completion of cell differentiation and frozen at −20°C for subsequent hormone analysis. Cell pellets were also collected in parallel and lysed in 250 µl RIPA buffer (Pierce, ThermoFisher Scientific, cat no: P18990). Protein extracts were diluted in PBS and quantified in duplicate using a Micro BCA Protein Assay Kit (ThermoFisher Scientific, cat no: 23235). Bovine serum albumin (BSA) of known protein concentrations was used as the standard curve to determine protein concentrations for the cell protein extracts. To quantify mCG secretion, a radioimmunoassay was used as previously described ^81, 82^. Samples were diluted in PBS and run in duplicate. The inter-assay variation (CV) was 6.0%. An enzyme immunoassay (EIA) (Cayman Chemical, cat no: 582601) was used to quantify progesterone secretion in cell conditioned media as previously published ^82^. Briefly, samples were diluted in enzyme-linked immunosorbent assay (ELISA) buffer and run in duplicate. A standard curve of known concentration was used to determine the sample concentrations. The minimum level of detection was 0.1ng/ml. For statistical analyses, samples below the lower limit of detection (0.1 ng/ml) were assigned 0.1ng/ml. Secretion was normalized to µg cell protein and for statistical analysis a log transformation was performed followed by a nonparametric Kruskal-Wallis test with Dunn’s correction was used to test significance (p < 0.05).

### Flow Cytometry

Cryopreserved cells were thawed at room temperature and washed twice with 10 ml of FACS buffer (PBS and 2% FBS) and filtered through a 100 µM filter (BD BioSciences, cat no: 352360). Cell concentration was determined by hemocytometer. Viability was assessed by staining with Ghost Red 780 (Tonbo Biosciences, cat no: 13-0865) dye for 30 min at 4°C. Cells were then washed and stained for surface cell markers, 25D3 (Mamu-AG) Alexa 647, and CD-56 PE-CF594 for 20 min at room temperature and then washed, see Supplemental Table S9 for flow cytometry antibody information. In house antibody 25D3 (Mamu-AG)^34^, was conjugated to Alexa 647 using the Biotium Mix-n-Stain R-PE(PE) Antibody Labeling kit (Biotium, cat no: 92298) following the manufacturers protocol. Cells were then fixed and permeabilized using the Foxp3 transcription factor staining buffer kit (eBioscience, cat no: 00-552-3-00) according to the manufacture’s guidelines. Fixation and washing were followed by intracellular staining using the following antibodies: Ki-67 Brilliant Violet 711, Cytokeratin 7/8 Brilliant Violet 421, Vimentin Alexa Flour 488. Intracellular staining was done for 30 min at 4°C. Unstained cells, control cell lines and compensation controls using cell lines or UltraComp eBeads (eBioscience, cat no: 01-2222-42) were prepared concurrently. All eight TSC cell lines, three EVT lines each generated with or without A83-01 supplementation and a rhesus fibroblast cell line were evaluated on a five laser BD LSRFortessa instrument. Positive staining was confirmed using positive and negative control cell lines. Results were analyzed using FlowJo v10.6 software (FlowJo, LLC). The gating strategy was determined based on unstained cells.

### DNA methylation by bisulfite sequencing

DNA was extracted from blastocyst stage embryos (n=3), rhesus fibroblast cell line (n=1), pri-CTB (n=1, rh090419) and TSCs (n=3 rh121118, rh052318, cy091318) using a FlexiGene DNA Kit (Qiagen, cat no: 51206) and quantified using a Nanodrop™ One Microvolume UV-Vis Spectrophotometer (ThermoFisher Scientific). DNA was bisulfite converted using an EZ DNA Methylation-Lightning™ Kit (Zymo Research Corporation, D5030T). PCR was performed to amplify the ELF5 and C19MC gene regions, see Supplemental Table S7 for bisulfite treated DNA primers. PCR products were purified using a QIAquick PCR Purification Kit (Qiagen, cat no: 28104) and were then cloned using a pGEM→-T Easy Vector System (Promega, cat no: A1380). For each sample, DNA was isolated from 30 clones using a QIAprep Spin Miniprep Kit (Qiagen, cat no: 27106). Clone DNA was then Sanger sequenced at the University of Wisconsin-Madison Biotechnology Center using the T7 primer of the vector. Methylation marks were identified for each CpG site within the gene region (ELF5 11 CpG sites; C19MC 34 CpG sites) each cell type. ELF5 methylation was analyzed by determine the percentage of clones with methylated CpGs at each site.

### Single-Cell Sequencing

Cryopreserved passage 2 and passage 10 cells were thawed and cultured for 72 h. TSCs were then lifted with TrypLE by incubation for 30 min at 37°C. An equal volume of DMEM/F12 was then added and cells were triturated by passage through various sizes of pipette tips. The cell suspension was pelleted, supernatant removed and resuspended in 1 ml DMEM/F-12 and cells were then strained through a 10 µm cell strainer followed by additional trituration. Single cell suspensions were then evaluated for viability and cell count by using a Countess II FL Automated Cell Counter (Invitrogen, ThermoFisher). Cells were then diluted to a concentration of 600 cells/µl for a target cell range of ∼5000 cells to be individually encapsulated into nanoliter-scale droplets containing reverse transcription reagents and uniquely barcoded Gel Bead-in EMulsions (GEMS). cDNA quality was assessed using an Agilent HS DNA chip and libraries were prepared using a Chromium Single Cell 3’ v2 kit (10x Genomics, USA). Libraries were quantified using a Qubit Fluorometer (Invitrogen) with a Qubit dsDNA HS kit (Invitrogen, ThermoFisher Scientific, cat no: Q33230) and evaluated for quality by sequencing on a MiSeq Nano platform (Illumina). Libraries were then sequenced across two lanes of a Novaseq 6000 SP flow cell (lllumina).

Single-cell RNA sequencing (scRNA-seq) data was analyzed by the UW Bioinformatics Resource Center. Sequencing data was demultiplexed using the Cell Ranger Single Cell Software Suite, mkfastq command wrapped around Illumina’s bcl2fastq. The MiSeq balancing run was quality controlled using calculations based on UMI-tools^83^. Samples’ libraries were balanced for the number of estimated reads per cell and run on an Illumina NovaSeq system. Cell Ranger software was then run to perform demultiplexing, alignment, filtering, barcode counting, UMI counting, and gene expression estimation for each sample according to the 10x Genomics documentation (https://support.10xgenomics.com/single-cell-gene-expression/software/pipelines/latest/what-is-cell-ranger). Reads were aligned to the rhesus macaque reference genome MacaM_v7.8.2. The gene expression estimates from each sample were then aggregated using Cellranger (cellranger aggr) to compare experimental groups with normalized sequencing-depth and expression data. A dimensionality reduction method, t-Stochastic Neighbor Embedding (t-SNE), was applied to plot hierarchical clusters of nearest neighbors in 2-dimensional space^84^. In addition, cell expression profiles are also represented by K-means clustering using Davies-Bouldin Index to identify the ideal number of preset number of clusters. For each t-SNE cluster, the differential expression algorithms are run to compare to all other cells, yielding a list of differentially expressed genes in that cluster compared to the rest of the sample. The up or down regulated significantly differentially expressed genes (p<0.05, >2-fold change expression) were imputed into the web-based tool, ENRICHR^85, 86^, to identify Human KEGG pathways and Gene Ontology Biological Processes altered by the gene set.

## Acknowledgements

The authors would like to thank the Wisconsin National Primate Research Center units including the Scientific Protocol Implementation, Veterinary Services, Colony Management and Pathology Services Units, especially Dr. Kevin Brunner and Michele Schotzko, for assisting in collection of macaque placentas. The authors also extend thanks to the University of Wisconsin-Madison Biotechnology Center Gene Expression Center and DNA Sequencing Facility for providing single cell library preparation and next generation sequencing services. We would also like to thank the University of Wisconsin-Madison Biotechnology Center Bioinformatics Resource Center, particularly Drs. Derek Pavelec and Mark Berres, for analysis of RNA-sequencing datasets. The authors would also like to acknowledge funding support by the following funding sources: P51 OD011106-54 to the Wisconsin National Primate Research Center; R01 AI132419-01A1, R01 AI107157-01A1, and R21 AI136012-01 to TGG; T32GM081061 and F31HD100057 to LNB; and K99 HD099154-01 to JKS. We also acknowledge support for the BD LSRFortessa flow cytometry instrument by the University of Wisconsin Carbone Cancer Center Support (NIH grant P30 CA014520).

## Author Contributions

TGG and JKS contributed to the conception and design of the study. JKS, MGM, LNB, GJW, BMD, KMK, MJB, LTK, KDM, and MRK collected and analyzed data as well as aided in the drafting of the manuscript. The publication’s contents are solely the responsibility of the authors and do not necessarily represent the official views of the NIH.

