## Supplemental Information for "Derivation of macaque trophoblast stem cells"

### Supplemental Figure Legends

**Supplemental Figure S1.** Time course of TSC and pri-ST cultures. A) Phase contrast images of TSC and pri-ST at 24, 48 and 72 h in culture. B) Immunocytochemistry for Mamu-AG (red), Ki-67 (pink, nuclear, white arrowhead), f-actin (green) and nuclear stain, DAPI (blue). Scale bars in all panels represent 100 µm.

**Supplemental Figure S2.** Immunocytochemistry IgG control images.

**Supplemental Figure S3.** Extravillous trophoblast (EVT) morphology by phase contrast imaging. EVTs were grow in the absence or presence of ALK-5 inhibitor A83-01. Representative images for EVTs derived from TSCs of rh090419 (A) and rh010319 (B). Culture on Matrigel versus col-IV for EVTs derived from TSCs of rh121118.

**Supplemental Figure S4.** Flow cytometry analysis of TSCs (A) and rhesus fibroblasts (B).

**Supplemental Figure S5.** Flow cytometry analysis of EVTs grown in the presence or absence of A83-01 on col IV (A) or grown on Matrigel (B).

**Supplemental Table S1.** Karyotype analysis of TSC lines

| TSC Line | Gestation Date | Gender | Passage | Result | Summary |
|---|---|---|---|---|---|
| ***rhesus*** | | | | | |
| rh052318 | 40 | XX | 35 | abnormal | 2 clones with translocations and inversion of chr 2, 3, 12. |
| | | | 10a | normal | 17 normal cells, 2 non-clonal aberrations |
| | | | 30a | abnormal | 10 normal cells, 4 cells trisomy chr 3, 6 cells non-clonal aberrations |
| rh121118 | 62 | XY | 13 | abnormal | 11 normal cells, 1 cell non-clonal aberrations, 3 abnormal clones: clone 1) 3 cells duplication chr 2, clone 2) 3 cells trisomy chr3, clone 3) 2 cells trisomy chr 3 |
| rh010319 | 58 | XY | 12 | normal | 18 normal cells, 2 cell non-clonal aberrations |
| rh012919 | 57 | XY | 7 | normal | 17 normal cells, 3 cells non-clonal aberrations |
| rh020119 | 74 | XX | 7 | normal | 17 normal cells, 3 non-clonal aberrations |
| rh090419 | 75 | XX | 8 | normal | 18 normal cells, 2 cell non-clonal aberrations |
| ***cynomolgus*** | | | | | |
| cy091318 | 50 | XY | 7 | abnormal | 11 normal cells, 9 non-clonal aberrations. 33% of metaphase cells were polyploidy |
| | | | 6a | abnormal | 3 cells analyzed, all were tetraploid |
| cy020519 | 57 | XX | 11 | abnormal | 11 normal cells, 5 cells trisomy chr 6, 4 cells non-clonal aberrations |

a: denotes a passage generated from a previous cryopreserved vial of cells; chr: chromosome

**Supplemental Table S2. RNA-seq analysis of rh121118 cells.** Gene expression is represented as transcripts per million.

| Gene Name | pri-CTB | TSC | pri-ST | ST-2D | ST-3D |
|---|---|---|---|---|---|
| KRT7 | 64.78 | 96.88 | 98.26 | 67.61 | 325.36 |
| KRT8 | 917.26 | 3347.73 | 2610.18 | 2300.72 | 4488.45 |
| KRT18 | 1069.15 | 4627.46 | 4898.16 | 2728.91 | 6067.56 |
| VIM | 313.73 | 2 | 377.77 | 0.49 | 0.27 |
| ELF5 | 6.44 | 0.09 | 0 | 0.65 | 2.48 |
| GATA2 | 49.1 | 52.69 | 9.86 | 244.39 | 193.52 |
| GATA3 | 13.72 | 30.47 | 10.62 | 28.06 | 24.54 |
| GATA4 | 0.2 | 0.01 | 0.03 | 0.1 | 0.07 |
| GATA6 | 0.75 | 17.29 | 0.43 | 2.97 | 3.24 |
| GCM1 | 560.54 | 2 | 579.13 | 307.66 | 278.22 |
| HAND1 | 0.1 | 5.22 | 0 | 19.45 | 4.37 |
| ID2 | 52.64 | 23.68 | 21.15 | 17.95 | 3.76 |
| TEAD4 | 15.79 | 64.04 | 2.78 | 19.93 | 10.35 |
| TFAP2A | 65.77 | 36.41 | 47.59 | 123.35 | 73.36 |
| TFAP2C | 43.79 | 96.05 | 93.05 | 105.12 | 132.61 |
| TP63 | 27.48 | 18.9 | 0.58 | 14.82 | 5.93 |
| PLAC1 | 28.29 | 14.34 | 54.85 | 18.72 | 40.13 |
| PLAC8 | 1308 | 64.76 | 2948.95 | 739.73 | 1759.73 |
| ERVFRD1 | 235.24 | 2.09 | 10.14 | 95.73 | 117.51 |
| CDH1 | 47.96 | 180.59 | 66.65 | 126.72 | 60.03 |
| ITGA1 | 215.9 | 16.71 | 456.42 | 291.16 | 263.43 |
| ITGA5 | 356.7 | 18.99 | 1076.73 | 467.91 | 542.53 |
| ITGA6 | 325.51 | 302.33 | 153.79 | 328.09 | 180.13 |
| MMP2 | 686.77 | 611.56 | 1831.27 | 4279.43 | 6015.36 |
| MMP9 | 2.14 | 0.09 | 41.93 | 0.08 | 1.38 |
| NCAM1 | 126.78 | 249.57 | 197.56 | 321.77 | 398.36 |
| SDC1 | 25.18 | 12.25 | 50.2 | 131.52 | 143.01 |
| CGA | 1227.43 | 334.23 | 585.2 | 25852.41 | 50906.56 |
| CGB8 | 0 | 0.38 | 0 | 911.61 | 1628.96 |
| CSH1 | 706.85 | 0.08 | 91.66 | 178.29 | 38.23 |
| CSH4 | 146.58 | 0 | 82.01 | 24.66 | 38.83 |
| CYP11A1 | 57.64 | 171.47 | 245.54 | 451.19 | 666.96 |
| CYP19A1 | 9.16 | 0.5 | 7.49 | 0.7 | 0.88 |
| ESRRB | 0 | 0 | 0 | 0.11 | 0 |

| | | | | | |
|---|---|---|---|---|---|
| GNRH1 | 2.13 | 1.36 | 3.87 | 1.24 | 2.08 |
| HSD17B1 | 0.21 | 9.02 | 5.23 | 11.56 | 11.01 |
| MAMUA | 327.58 | 261.09 | 1115.74 | 631.94 | 632.46 |
| MAMUE | 99.55 | 199.96 | 299.25 | 224 | 312.51 |
| MAMUAG | 1061.53 | 342.37 | 3825.23 | 2181.64 | 2397.55 |

**Supplemental Table S3.** miRNA expression in pregnancy-associated miRNA clusters represented as transcripts per million

| | pri-CTB | TSC | pri-ST | ST-2D | ST-3D | EVT | fibroblast |
|---|---|---|---|---|---|---|---|
| **miR-371-3 cluster** | | | | | | | |
| miR-371-3p | 0 | 1 | 0 | 1 | 0 | 1 | 0 |
| miR-371-5p | 11 | 884 | 23 | 124 | 56 | 9 | 0 |
| miR-372-3p | 6 | 207 | 20 | 52 | 25 | 3 | 0 |
| miR-372-5p | 0 | 3 | 0 | 0 | 0 | 0 | 0 |
| miR-373 | 6 | 147 | 6 | 25 | 18 | 1 | 0 |
| **C19MC** | | | | | | | |
| miR-512-3p | 323 | 134 | 463 | 200 | 476 | 64 | 0 |
| miR-512-5p | 2706 | 3064 | 3329 | 1901 | 9167 | 1473 | 0 |
| miR-1323-3p | 0 | 0 | 3 | 0 | 2 | 0 | 0 |
| miR-1323-5p | 7263 | 9535 | 17798 | 10796 | 18964 | 7026 | 5 |
| miR-498-5p | 2548 | 2008 | 3321 | 1960 | 5295 | 1016 | 0 |
| miR-519e | 23361 | 23081 | 38525 | 24422 | 51396 | 8382 | 0 |
| miR-1283 | 583 | 320 | 492 | 338 | 352 | 80 | 0 |
| miR-520a-3p | 1455 | 1209 | 1882 | 1783 | 3497 | 465 | 0 |
| miR-520a-5p | 1863 | 939 | 1974 | 2228 | 1294 | 446 | 0 |
| miR-526b | 5524 | 2766 | 9295 | 3723 | 5678 | 1284 | 0 |
| miR-525 | 392 | 141 | 436 | 363 | 349 | 81 | 0 |
| miR-523a | 492 | 230 | 601 | 316 | 500 | 94 | 0 |
| miR-518f | 10919 | 11651 | 15307 | 13635 | 15787 | 4859 | 0 |
| miR-519a-3p | 46 | 47 | 8 | 69 | 58 | 3 | 0 |
| miR-519a-5p | 484 | 867 | 805 | 781 | 1355 | 457 | 0 |
| miR-518b | 7215 | 4904 | 8910 | 4982 | 8068 | 1942 | 1 |
| miR-518c-5p | 43 | 41 | 90 | 40 | 66 | 17 | 0 |
| miR-518c-3p | 4095 | 1256 | 3532 | 2119 | 1785 | 318 | 0 |
| miR-524-5p | 668 | 433 | 780 | 783 | 1414 | 160 | 0 |
| miR-524-3p | 28 | 24 | 37 | 32 | 38 | 1 | 0 |
| miR-517a | 2598 | 1148 | 2043 | 1701 | 2054 | 276 | 0 |
| miR-519d | 1036 | 472 | 997 | 751 | 782 | 130 | 0 |
| miR-518a-3p | 1632 | 1642 | 3748 | 2207 | 6296 | 523 | 0 |
| miR-518a-5p | 437 | 361 | 398 | 369 | 669 | 64 | 0 |
| miR-520g-3p | 8 | 1 | 5 | 5 | 4 | 0 | 0 |
| miR-518d-3p | 552 | 573 | 757 | 795 | 1035 | 131 | 0 |
| miR-518d-5p | 6598 | 7371 | 17889 | 8850 | 17281 | 4809 | 1 |
| miR-523b | 381 | 289 | 364 | 421 | 454 | 87 | 0 |
| miR-516b | 2116 | 1309 | 2990 | 2335 | 2943 | 380 | 0 |

| | | | | | | | |
|---|---|---|---|---|---|---|---|
| miR-518a-3p | 1632 | 1642 | 3748 | 2207 | 6296 | 523 | 0 |
| miR-518a-5p | 437 | 361 | 398 | 369 | 669 | 64 | 0 |
| miR-517c | 4474 | 5917 | 8006 | 7528 | 11482 | 2061 | 0 |
| miR-519b | 76 | 62 | 154 | 86 | 182 | 35 | 0 |
| miR-521 | 11 | 8 | 16 | 6 | 27 | 4 | 0 |
| miR-518e | 39 | 50 | 110 | 97 | 213 | 32 | 0 |
| miR-518a-3p | 1632 | 1642 | 3748 | 2207 | 6296 | 523 | 0 |
| miR-518a-5p | 437 | 361 | 398 | 369 | 669 | 64 | 0 |
| miR-516a-3p | 1085 | 1371 | 2165 | 1906 | 2489 | 531 | 0 |
| miR-516a-5p | 21483 | 10598 | 43883 | 15764 | 21448 | 4654 | 1 |
| **C14MC** | | | | | | | |
| miR-770-5p | 0 | 0 | 0 | 0 | 0 | 0 | 1 |
| miR-493-3p | 12 | 0 | 0 | 0 | 0 | 0 | 17 |
| miR-493-5p | 108 | 0 | 31 | 0 | 0 | 0 | 293 |
| miR-337-3p | 8 | 0 | 3 | 0 | 0 | 0 | 26 |
| miR-337-5p | 10 | 0 | 0 | 0 | 0 | 0 | 10 |
| miR-665 | 1 | 0 | 0 | 0 | 0 | 0 | 3 |
| miR-431 | 529 | 0 | 69 | 0 | 1 | 0 | 253 |
| miR-433-3p | 6 | 0 | 2 | 0 | 0 | 0 | 25 |
| miR-433-5p | 1 | 0 | 0 | 0 | 0 | 0 | 0 |
| miR-127-3p | 1571 | 0 | 203 | 0 | 0 | 0 | 2193 |
| miR-127-5p | 4 | 0 | 1 | 0 | 0 | 0 | 0 |
| miR-432-3p | 0 | 0 | 0 | 0 | 0 | 0 | 2 |
| miR-432-5p | 775 | 0 | 167 | 0 | 0 | 1 | 3776 |
| miR-136 | 5 | 0 | 0 | 0 | 0 | 0 | 0 |
| miR-370-3p | 65 | 0 | 3 | 0 | 0 | 0 | 173 |
| miR-370-5p | 3 | 0 | 1 | 0 | 0 | 0 | 16 |
| miR-379-3p | 5 | 0 | 0 | 0 | 0 | 0 | 18 |
| miR-379-5p | 217 | 0 | 22 | 0 | 0 | 0 | 309 |
| miR-411-3p | 2 | 0 | 0 | 0 | 0 | 0 | 11 |
| miR-411-5p | 363 | 0 | 19 | 0 | 0 | 0 | 280 |
| miR-299-3p | 88 | 0 | 7 | 0 | 0 | 0 | 50 |
| miR-299-5p | 427 | 0 | 82 | 0 | 2 | 3 | 992 |
| miR-380-3p | 0 | 0 | 0 | 0 | 0 | 0 | 4 |
| miR-380-5p | 0 | 0 | 0 | 0 | 0 | 0 | 2 |
| miR-323a-3p | 35 | 0 | 4 | 0 | 0 | 0 | 145 |
| miR-323a-5p | 0 | 0 | 0 | 0 | 0 | 0 | 0 |
| miR-758-3p | 2 | 0 | 0 | 0 | 0 | 0 | 5 |
| miR-758-5p | 1 | 0 | 0 | 0 | 0 | 0 | 0 |

| | | | | | | | |
|---|---|---|---|---|---|---|---|
| miR-329-3p | 13 | 0 | 1 | 0 | 0 | 0 | 56 |
| miR-329-1-5p | 0 | 0 | 0 | 0 | 0 | 0 | 2 |
| miR-329-2-5p | 0 | 0 | 0 | 0 | 0 | 0 | 0 |
| miR-494-3p | 15 | 0 | 4 | 0 | 0 | 0 | 50 |
| miR-494-5p | 0 | 0 | 0 | 0 | 0 | 0 | 1 |
| miR-543-3p | 13 | 0 | 2 | 0 | 0 | 0 | 38 |
| miR-543-5p | 4 | 0 | 0 | 0 | 0 | 0 | 13 |
| miR-495-3p | 10 | 0 | 0 | 0 | 0 | 0 | 11 |
| miR-495-5p | 5 | 0 | 1 | 0 | 0 | 0 | 9 |
| miR-376c-3p | 121 | 0 | 7 | 0 | 0 | 0 | 52 |
| miR-376c-5p | 6 | 0 | 0 | 0 | 0 | 0 | 1 |
| miR-376a-3p | 60 | 0 | 6 | 1 | 0 | 0 | 46 |
| miR-376a-1-5p | 3 | 0 | 1 | 0 | 0 | 0 | 0 |
| miR-376a-2-5p | 1 | 0 | 0 | 0 | 0 | 0 | 0 |
| miR-654-3p | 386 | 0 | 57 | 0 | 0 | 2 | 1631 |
| miR-654-5p | 0 | 0 | 0 | 0 | 0 | 0 | 0 |
| miR-376b-3p | 6 | 0 | 2 | 0 | 0 | 0 | 2 |
| miR-376b-5p | 3 | 0 | 0 | 0 | 0 | 0 | 1 |
| miR-376a-1-5p | 3 | 0 | 1 | 0 | 0 | 0 | 0 |
| miR-376a-2-5p | 1 | 0 | 0 | 0 | 0 | 0 | 0 |
| miR-376a-3p | 60 | 0 | 6 | 1 | 0 | 0 | 46 |
| miR-1185-3p | 3 | 0 | 1 | 0 | 0 | 0 | 10 |
| miR-1185-5p | 8 | 0 | 0 | 0 | 0 | 0 | 2 |
| miR-381-3p | 77 | 0 | 9 | 0 | 0 | 1 | 88 |
| miR-381-5p | 0 | 0 | 0 | 0 | 0 | 0 | 1 |
| miR-487b-3p | 110 | 0 | 18 | 0 | 0 | 0 | 183 |
| miR-487b-5p | 4 | 0 | 0 | 0 | 0 | 0 | 30 |
| miR-539 | 0 | 0 | 0 | 0 | 0 | 0 | 0 |
| miR-889-3p | 53 | 0 | 4 | 0 | 0 | 0 | 44 |
| miR-889-5p | 0 | 0 | 0 | 0 | 0 | 0 | 0 |
| miR-544 | 0 | 0 | 0 | 0 | 0 | 0 | 0 |
| miR-487a | 0 | 0 | 0 | 0 | 0 | 0 | 0 |
| miR-382-3p | 27 | 0 | 5 | 0 | 0 | 0 | 69 |
| miR-382-5p | 371 | 0 | 76 | 0 | 0 | 0 | 1164 |
| miR-134-3p | 0 | 0 | 0 | 0 | 0 | 0 | 1 |
| miR-134-5p | 95 | 0 | 12 | 0 | 0 | 2 | 544 |
| miR-668 | 4 | 0 | 1 | 0 | 0 | 0 | 9 |
| miR-485-3p | 46 | 0 | 14 | 0 | 1 | 0 | 1102 |

| | | | | | | | |
|---|---|---|---|---|---|---|---|
| miR-485-5p | 32 | 0 | 2 | 0 | 0 | 0 | 201 |
| miR-323b-3p | 28 | 0 | 6 | 0 | 0 | 0 | 279 |
| miR-323b-5p | 0 | 0 | 0 | 0 | 0 | 0 | 0 |
| miR-154-3p | 2 | 0 | 0 | 0 | 0 | 0 | 6 |
| miR-154-5p | 48 | 0 | 8 | 0 | 0 | 0 | 54 |
| miR-496 | 4 | 0 | 0 | 0 | 0 | 0 | 7 |
| miR-337-3p | 8 | 0 | 3 | 0 | 0 | 0 | 26 |
| miR-337-5p | 10 | 0 | 0 | 0 | 0 | 0 | 10 |
| miR-541-3p | 0 | 0 | 0 | 0 | 0 | 0 | 0 |
| miR-541-5p | 0 | 0 | 0 | 0 | 0 | 0 | 10 |
| miR-409-3p | 532 | 0 | 73 | 0 | 0 | 1 | 2165 |
| miR-409-5p | 34 | 0 | 5 | 0 | 0 | 0 | 115 |
| miR-412-3p | 0 | 0 | 0 | 0 | 0 | 0 | 1 |
| miR-412-5p | 33 | 0 | 5 | 0 | 0 | 0 | 16 |
| miR-369-3p | 78 | 0 | 1 | 0 | 0 | 0 | 45 |
| miR-369-5p | 166 | 0 | 15 | 0 | 0 | 0 | 335 |
| miR-410-3p | 2 | 0 | 0 | 0 | 0 | 0 | 3 |
| mIR-410-5p | 0 | 0 | 0 | 0 | 0 | 0 | 13 |
| miR-656-3p | 3 | 0 | 0 | 0 | 0 | 0 | 11 |
| miR-656-5p | 0 | 0 | 0 | 0 | 0 | 0 | 1 |
| miR-1247-3p | 7 | 0 | 0 | 0 | 0 | 0 | 0 |
| miR-1247-5p | 118 | 0 | 14 | 0 | 0 | 0 | 0 |

**Supplemental Table S4.** Differentially expressed genes between passage 10 TSC clusters identified by scRNA-Seq t-SNE analysis. Parenthesis indicate number of genes either significantly down or upregulated ($p < 0.05$) and greater than 2-fold change in expression

| **Cluster 1** | **Cluster 2** | **Cluster 4** | **Cluster 5** | **Cluster 6** | **Cluster 7** | **Cluster 9** |
|---|---|---|---|---|---|---|
| **Up (12)** | **Down (7)** | **Down (1)** | **Down (9)** | **Down (1)** | **Up (6)** | **Down (2)** |
| TOP2A | TOP2A | TOP2A | TOP2A | TOP2A | TOP2A | EDA2R |
| PRC1 | PRC1 | | PRC1 | | UBE2S | IRS2 |
| CHAC1 | H2AFX | | CHAC1 | | PRC1 | |
| PRRG4 | UBE2S | | BIRC5 | | BIRC5 | |
| EDA2R | BIRC5 | | WEE1 | | ARL6IP1 | |
| RHNO1 | CKAP5 | | UBE2S | | CKAP5 | |
| SUN2 | ARL6IP1 | | IRS2 | | | |
| UBE2S | | | SLC7A1 | | **Down (2)** | |
| MYADM | | | H2AFX | | IRS2 | |
| WEE1 | | | | | THBS1 | |
| H2AFX | | | | | | |
| BIRC5 | | | | | | |

**Supplemental Table S5.** Differentially expressed genes between TSC clusters identified by scRNA-Seq K-means analysis of p2 and p10 cells combined. Parenthesis indicate number of genes either significantly down or upregulated (p <0.05) and greater than 2-fold change in expression

| **Cluster 1** | **Cluster 2** |
|---|---|
| **Down (187)** | **Up (171)** |
| APOE | NUCB2 |
| NUCB2 | UBB |
| UBB | TECR |
| TECR | PMEL |
| PMEL | UBC |
| PLTP | APOE |
| UBC | MRPS31 |
| TUBA3C | PRDX4 |
| PRDX4 | PLTP |
| MRPS31 | LDHA |
| LDHA | TUBA3C |
| PRDX2 | PSMB3 |
| PSMB3 | ANXA2 |
| CTNNBL1 | PRDX2 |
| ANXA2 | CTNNBL1 |
| NSDHL | CDK1 |
| CDK1 | NSDHL |
| CKB | EIF4A3 |
| DRG1 | CKB |
| EIF4A3 | DRG1 |
| BLVRB | PSME2 |
| RBM42 | TYMS |
| PSME2 | PGK1 |
| SUGT1 | WDR61 |
| ISYNA1 | SUGT1 |
| PGK1 | BLVRB |
| TYMS | PSMC4 |
| PSMC4 | DDX39A |
| DDX39A | RBM42 |
| WDR61 | CD9 |
| KPNA2 | KPNA2 |
| CD9 | PSMD12 |

| | |
|---|---|
| PSMD12 | ISYNA1 |
| FOLR1 | PNP |
| C22orf28 | LDHB |
| LDHB | GHITM |
| PNP | ATP5B |
| GHITM | C22orf28 |
| ATP5B | FOLR1 |
| RUVBL2 | RUVBL2 |
| KRT18 | UFD1L |
| UFD1L | PSMA6 |
| PSMC5 | CLDN6 |
| CLDN6 | KRT18 |
| PDIA3 | TK1 |
| FAM50A | PDIA3 |
| PSMA6 | PRMT1 |
| PSMD7 | PSMC5 |
| PRMT1 | PSMD7 |
| PKM | NUDT5 |
| ACP5 | RAB11A |
| TK1 | UQCRC2 |
| PLD3 | PKM |
| RAB11A | FAM50A |
| NUDT5 | TUBA1B |
| UQCRC2 | KRT7 |
| PRPF19 | ITM2B |
| KRT7 | ACP5 |
| ITM2B | PRPF19 |
| AURKB | RDM1 |
| TUBA1B | MYL12B |
| RDM1 | AURKB |
| CTSA | AHSA1 |
| PSMC3 | LGALS3 |
| MYL12B | PSMC1 |
| AHSA1 | UBE2C |
| MVD | MVD |
| PSMC1 | PYGL |
| GRN | PSMA3 |
| UBE2C | NDUFA9 |
| LGALS3 | PA2G4 |

| | |
|---|---|
| XRCC6 | PLK1 |
| PYGL | PSMC3 |
| PA2G4 | MYL12A |
| WDR18 | XRCC6 |
| PLK1 | WDR18 |
| TMEM205 | CTSA |
| PSMA3 | GOT1 |
| NDUFA9 | PLD3 |
| HMOX1 | EML2 |
| GOT1 | CDC123 |
| MYL12A | RNASEH2A |
| EML2 | HMOX1 |
| PPP2R1A | TMEM205 |
| RNASEH2A | ATP5C1 |
| PPIB | PPP2R1A |
| NDUFV1 | SNRPB2 |
| COMMD4 | GRN |
| CDC123 | NDUFV1 |
| ATP6V0D1 | COMMD4 |
| CLDN7 | TPI1 |
| SNRPB2 | MAGEA4 |
| HSP90B1 | PPIB |
| PVRL2 | ATP6V0D1 |
| IRS2 | CLDN7 |
| TPI1 | MCTS1 |
| ADRM1 | TUBG1 |
| ATP5C1 | EIF2S1 |
| MCTS1 | MRPL46 |
| HEXA | ADRM1 |
| TUBG1 | HSP90B1 |
| MAGEA4 | MPV17L2 |
| EIF2S1 | RSU1 |
| GPS1 | ACAT1 |
| SLC16A3 | ATP5H |
| CYBA | SAR1A |
| ETFB | ORMDL2 |
| CD151 | CD151 |
| NOSIP | AP1M2 |
| MPV17L2 | GPS1 |

| | |
|---|---|
| RSU1 | SLC16A3 |
| MRPL46 | MYL6 |
| CDC37 | NSMCE1 |
| ORMDL2 | TALDO1 |
| ACAT1 | ESD |
| ATP5H | KRR1 |
| SAR1A | CDC37 |
| NSMCE1 | POLR2C |
| AP1M2 | HEXA |
| PHB | ANP32A |
| MYL6 | PSMB6 |
| HSD17B14 | PHB |
| PSMD3 | NOSIP |
| SF3B2 | PVRL2 |
| ANP32A | SF3B2 |
| GSTP1 | PSME1 |
| TALDO1 | GSTP1 |
| PSMB6 | HSD17B14 |
| POLR2C | ETFB |
| TIMM50 | EXOSC8 |
| ESD | TIMM50 |
| PAGE4 | TMBIM6 |
| TAC3 | NLRP2 |
| KRR1 | DPM1 |
| NLRP2 | ARPC3 |
| TMBIM6 | NDUFS3 |
| BCAP31 | BCAP31 |
| NDUFS3 | MTCH2 |
| FUS | CYBA |
| PELP1 | PSMD3 |
| NEMF | FUS |
| PSME1 | VGLL1 |
| PRSS8 | PAGE2B |
| PAGE2B | MAGOHB |
| EXOSC8 | PCNA |
| VGLL1 | RSL1D1 |
| DPM1 | TRAPPC4 |
| MFGE8 | TSG101 |
| PCNA | MRPL48 |

| | |
|---|---|
| ARPC3 | TAC3 |
| MVB12A | HSD17B10 |
| RSL1D1 | PAGE4 |
| VPS25 | NEMF |
| TSG101 | VPS25 |
| TUFM | SPCS2 |
| MTCH2 | HNRNPH3 |
| RBM25 | TXNL1 |
| MRPL48 | CHMP4A |
| HNRNPH3 | GLRX3 |
| HSD17B10 | PELP1 |
| SPCS2 | MVB12A |
| MAGOHB | PRSS8 |
| CHMP4A | RAN |
| TRAPPC4 | TUFM |
| BSG | STIP1 |
| TXNL1 | COMMD3 |
| ILK | ACAA2 |
| AMDHD2 | EIF4A1 |
| GLRX3 | DERA |
| TOMM40 | MFGE8 |
| STIP1 | RBM25 |
| ATP6AP1 | |
| MRPL38 | **Down (1)** |
| RAN | IRS2 |
| TUBA1C | |
| POLR2E | |
| MTHFD1 | |
| TIMM23 | |
| DERA | |
| DAD1 | |
| EIF4A1 | |
| SNRNP70 | |
| ACAA2 | |
| COMMD3 | |
| STX8 | |
| AKAP8L | |
| JUNB | |

**Supplemental Table S6.** Differentially expressed genes between TSC clusters segregated by the LMO algorithm applied to scRNA-seq gene expression levels of combined p2 and p10 cells. Parenthesis indicate number of genes either significantly down or upregulated (p <0.05) and greater than 2-fold change in expression

| **Cluster 1** | **Cluster 2** | **Cluster 3** | **Cluster 4** | **Cluster 5** | **Cluster 6** | **Cluster 8** | **Cluster 9** |
|---|---|---|---|---|---|---|---|
| **Up (3)** | **Down (75)** | **Up (13)** | **Up (1)** | **Up (2)** | **Down (8)** | **Up (95)** | **Up (51)** |
| APOE | CDK1 | WEE1 | PLEKHF1 | TOP2A | PLK1 | CCNB2 | MCM5 |
| PMEL | UBE2C | THBS1 | | GTSE1 | UBE2C | PLK1 | TK1 |
| FOLR1 | TOP2A | BMP7 | **Down (24)** | | APOE | CDCA3 | RAD51 |
| | PRC1 | FADS2 | CDK1 | **Down (28)** | CCNB2 | UBE2C | PCNA |
| **Down (19)** | GTSE1 | MT1E | UBE2C | PMEL | UBB | KPNA2 | FEN1 |
| TOP2A | CDCA3 | MYADM | TOP2A | TAC3 | PMEL | AURKB | PRIM1 |
| GTSE1 | TK1 | EDA2R | AURKB | APOE | CDCA3 | CDK1 | GOT1 |
| UBE2C | KPNA2 | CGB | CDCA3 | PLTP | UBC | TUBA3C | ORC6 |
| PRC1 | PLK1 | | PRC1 | ACP5 | | DLGAP5 | LDHA |
| CDK1 | RAD51 | **Down (40)** | TK1 | PAGE2 | | NUP37 | NUCB2 |
| CDCA3 | AURKB | APOE | GTSE1 | CD9 | | UBB | MPV17L2 |
| AURKB | UBB | TECR | PLK1 | PSMB3 | | TUBA1B | PA2G4 |
| PLK1 | TPX2 | PSMB3 | KPNA2 | PRDX4 | | DDX39A | NSDHL |
| TPX2 | KIF20B | PSME1 | TUBA3C | NUCB2 | | KIF20B | MVD |
| BIRC5 | MCM5 | LDHB | KIF20B | PRDX2 | | RBM42 | KRT7 |
| CASC5 | CTNNBL1 | BLVRB | RAD51 | BMP7 | | MRPS31 | TECR |
| UBE2S | APOE | PRDX2 | NUP37 | CLDN6 | | TYMS | PLTP |
| KIF20B | NUCB2 | PRDX4 | MCM5 | UBB | | ARL6IP1 | CTNNBL1 |
| H2AFX | PMEL | NUCB2 | TECR | ATP5B | | PSMD12 | UBC |
| MCM10 | UBC | FOLR1 | APOE | PSME2 | | RDM1 | CLDN6 |
| BRCA2 | GOT1 | ANXA2 | NUCB2 | BLVRB | | UBC | PRDX4 |
| TK1 | CKB | MRPS31 | UBB | TECR | | NUCB2 | KRT18 |
| CHAF1A | TUBA3C | UBB | UBC | MAGEH1 | | PRC1 | LGALS3 |
| RHNO1 | NUP85 | SLC25A5 | PKMYT1 | ANXA2 | | CTNNBL1 | CLDN7 |
| | CLDN6 | PSME2 | ANXA2 | PNP | | TUBA1C | PMEL |
| | PCNA | PSMC4 | TPX2 | UBC | | CDKN3 | APOE |
| | BIRC5 | PSMA3 | EIF4A3 | MCM5 | | UBE2S | PKM |
| | DRG1 | TUBA3C | | FOLR1 | | C22orf28 | CD9 |
| | LDHA | KRT19 | | MRPS31 | | DRG1 | CKB |

| | | | | | | | |
|---|---|---|---|---|---|---|---|
| | TAC3 | PMEL | | LDHA | | TBL3 | PPAN |
| | PLTP | SUGT1 | | CKB | | PSMD7 | PRDX2 |
| | ORC6 | ESD | | ISYNA1 | | TOP2A | PGK1 |
| | FEN1 | UBC | | | | RUVBL2 | EIF4A3 |
| | BMP7 | KRT18 | | | | LDHA | MCM10 |
| | PSMD7 | ISYNA1 | | | | SUGT1 | LDHB |
| | ATP5B | NSDHL | | | | PSMB3 | GINS2 |
| | PRIM1 | PSMC5 | | | | RAD51 | CHAF1A |
| | RANGAP1 | WDR61 | | | | NUP85 | DRG1 |
| | ARL6IP1 | RPN2 | | | | AHSA1 | NUDT5 |
| | NUP37 | RNASEH2A | | | | CKB | PRPF19 |
| | PKM | RBM42 | | | | SAAL1 | MPHOSPH8 |
| | UBE2S | ACAT1 | | | | MARS | NUP85 |
| | PSMD12 | EIF4A3 | | | | TECR | BLVRB |
| | ANXA2 | PLTP | | | | CKLF | UBB |
| | MRPS31 | KRT7 | | | | NSDHL | PNP |
| | PKMYT1 | TUFM | | | | PRDX4 | FOLR1 |
| | RUVBL2 | RUVBL2 | | | | PSMC4 | ANXA2 |
| | CENPJ | MCTS1 | | | | RNASEH2A | RAB11A |
| | DLGAP5 | NUP37 | | | | BIRC5 | HSP90B1 |
| | H2AFX | PSMA6 | | | | PGK1 | PYGL |
| | MCM10 | | | | | FAM50A | TUBA3C |
| | CASC5 | | | | | TPX2 | |
| | GHITM | | | | | CARS | Down (1) |
| | TECR | | | | | EIF4A3 | EDA2R |
| | PRDX4 | | | | | TK1 | |
| | NSDHL | | | | | PSMA6 | |
| | ACP5 | | | | | KRR1 | |
| | LGMN | | | | | PSMC5 | |
| | NLRP2 | | | | | PLTP | |
| | EIF4A3 | | | | | PSMC1 | |
| | CD9 | | | | | CIAPIN1 | |
| | TYMS | | | | | PRDX2 | |
| | PSMC3 | | | | | PMEL | |
| | C22orf28 | | | | | MYL12A | |
| | PDIA3 | | | | | GTSE1 | |
| | TUBA1B | | | | | NUDT5 | |
| | CTSA | | | | | FUS | |
| | TBL3 | | | | | PRMT1 | |

| | | | | | | |
|---|---|---|---|---|---|---|
| | DDX39A | | | | | GHITM |
| | MPV17L2 | | | | | PSMD3 |
| | SERTAD1 | | | | | EIF2S1 |
| | PLD3 | | | | | POLR2C |
| | PSMC4 | | | | | PSMC3 |
| | FAM50A | | | | | XRCC6 |
| | DHCR7 | | | | | UQCRC2 |
| | | | | | | WDR18 |
| | | | | | | SNRPB2 |
| | | | | | | HMGB1 |
| | | | | | | PSME2 |
| | | | | | | UFD1L |
| | | | | | | RAB11A |
| | | | | | | SARS2 |
| | | | | | | MAGOHB |
| | | | | | | STIP1 |
| | | | | | | PRPF19 |
| | | | | | | TUBG1 |
| | | | | | | APOE |
| | | | | | | SNRNP70 |
| | | | | | | LDHB |
| | | | | | | ANXA2 |
| | | | | | | GOT1 |
| | | | | | | PNP |
| | | | | | | WDR61 |
| | | | | | | RPN2 |
| | | | | | | FTHL17 |
| | | | | | | **Down (6)** |
| | | | | | | IRS2 |
| | | | | | | CRISPLD2 |
| | | | | | | DUSP5 |
| | | | | | | SLC7A8 |
| | | | | | | FOSB |
| | | | | | | MARVELD1 |
