## Supplementary figures and images for "Derivation of macaque trophoblast stem cells"

### Supplemental Figure S1

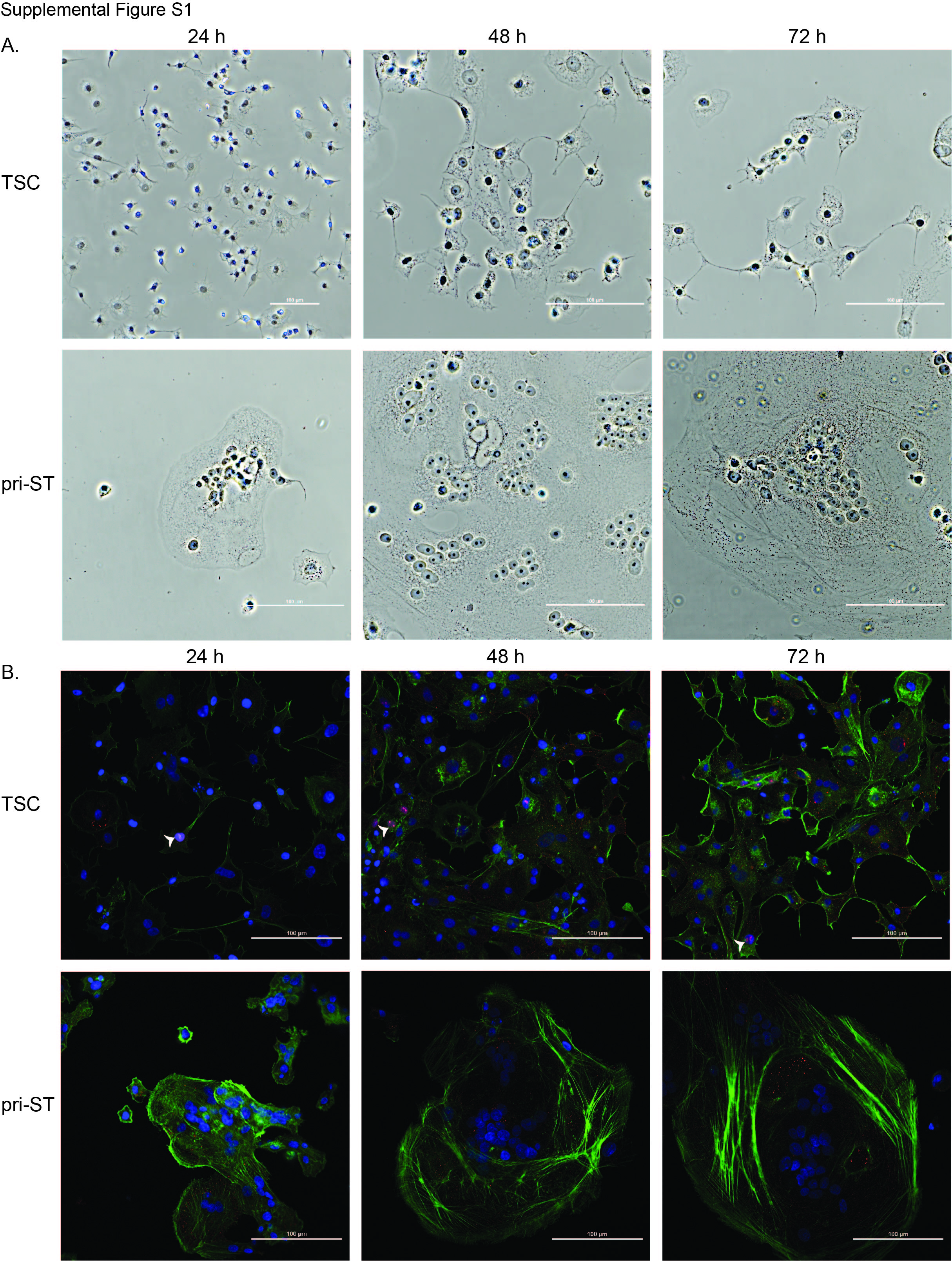

### Supplemental Figure S2

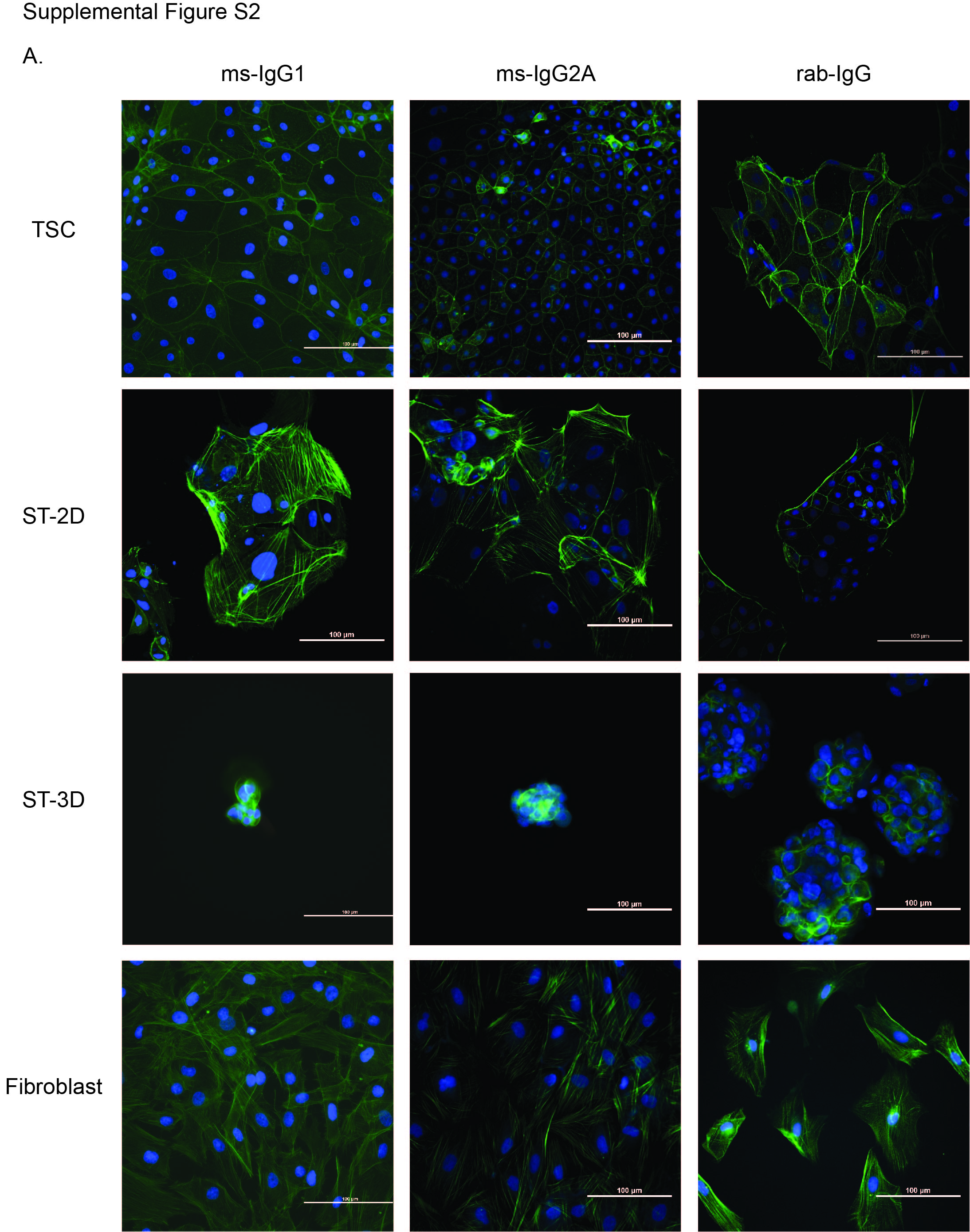

### Supplemental Figure S3

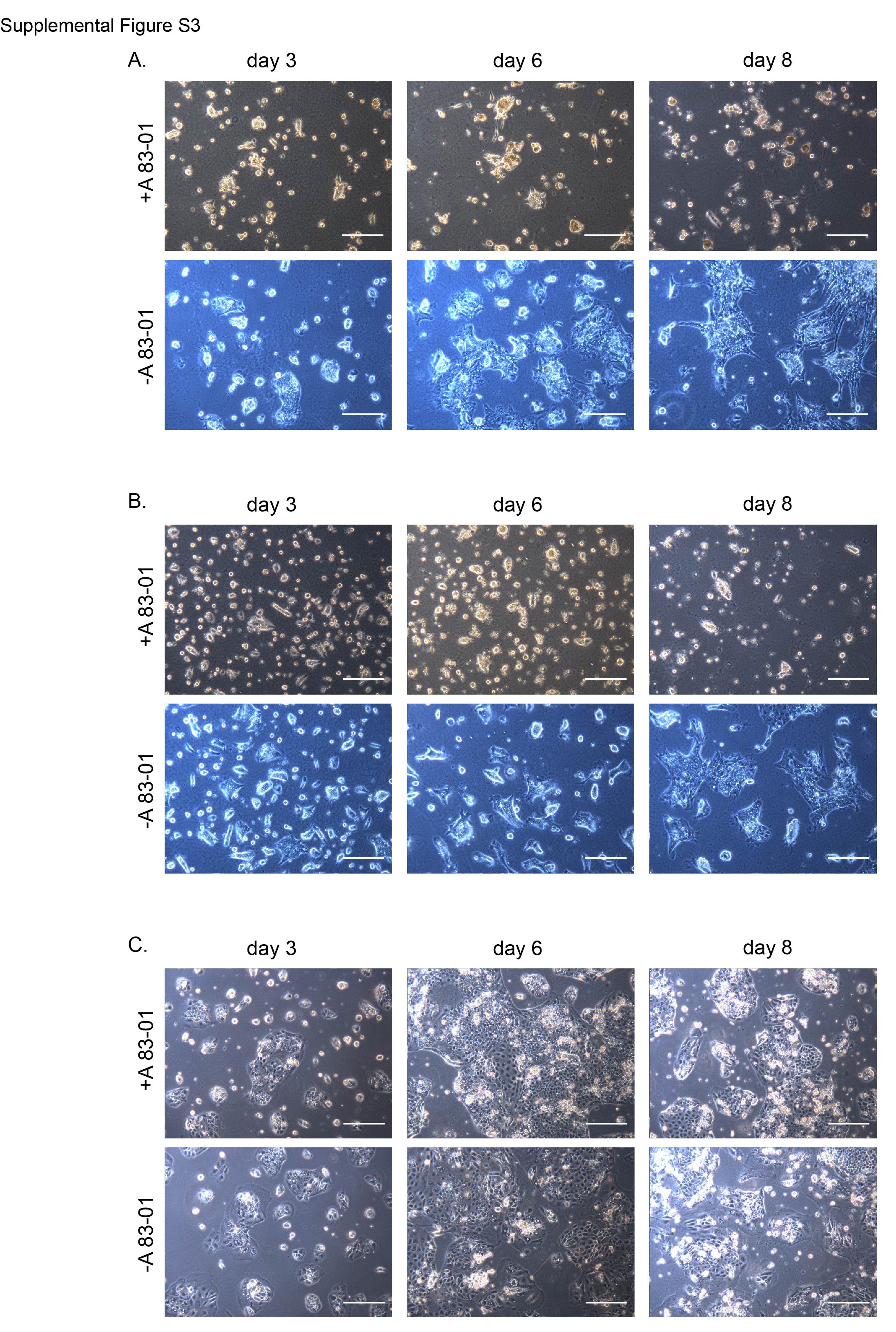

### Supplemental Figure S4

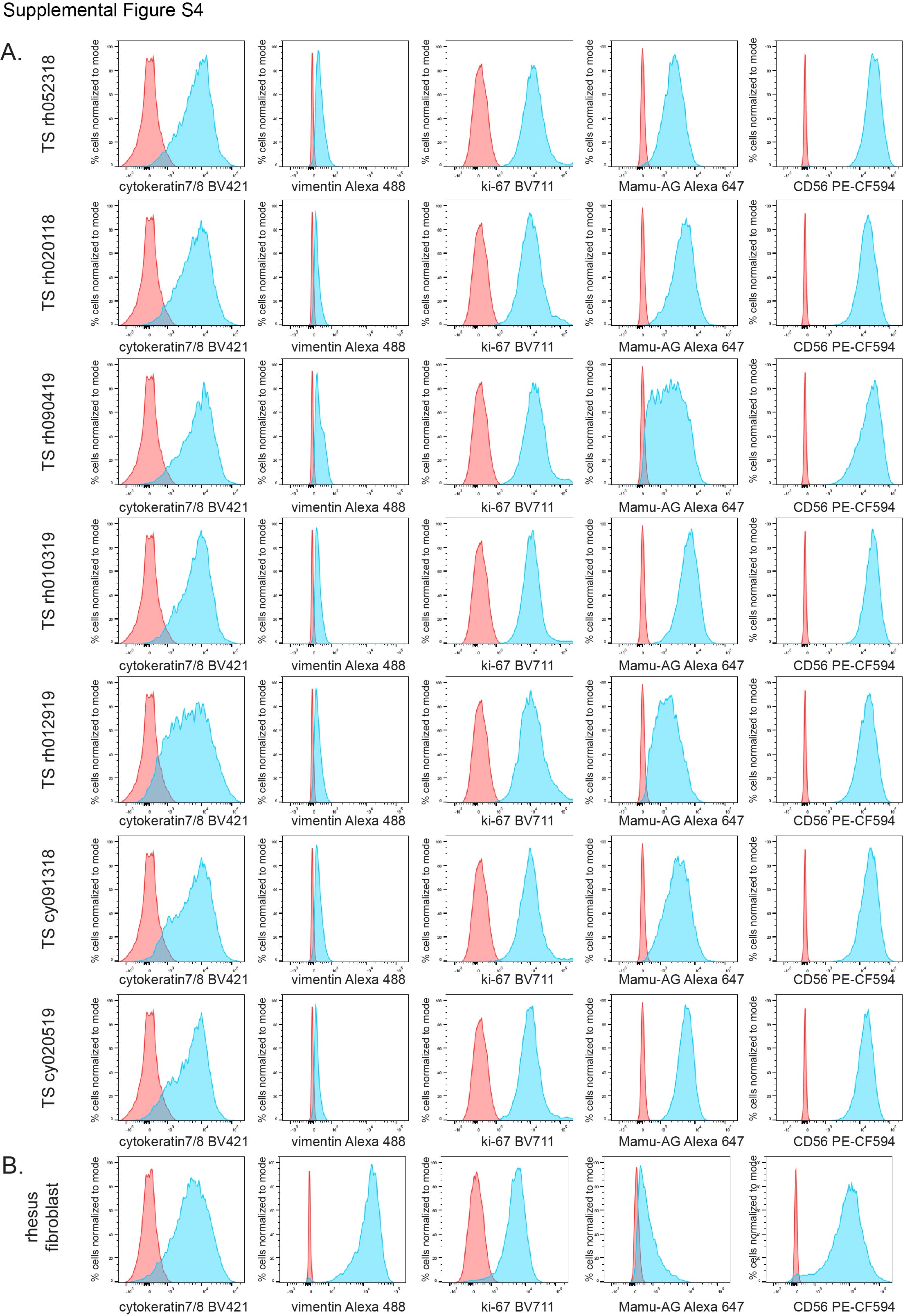

### Supplemental Figure S5

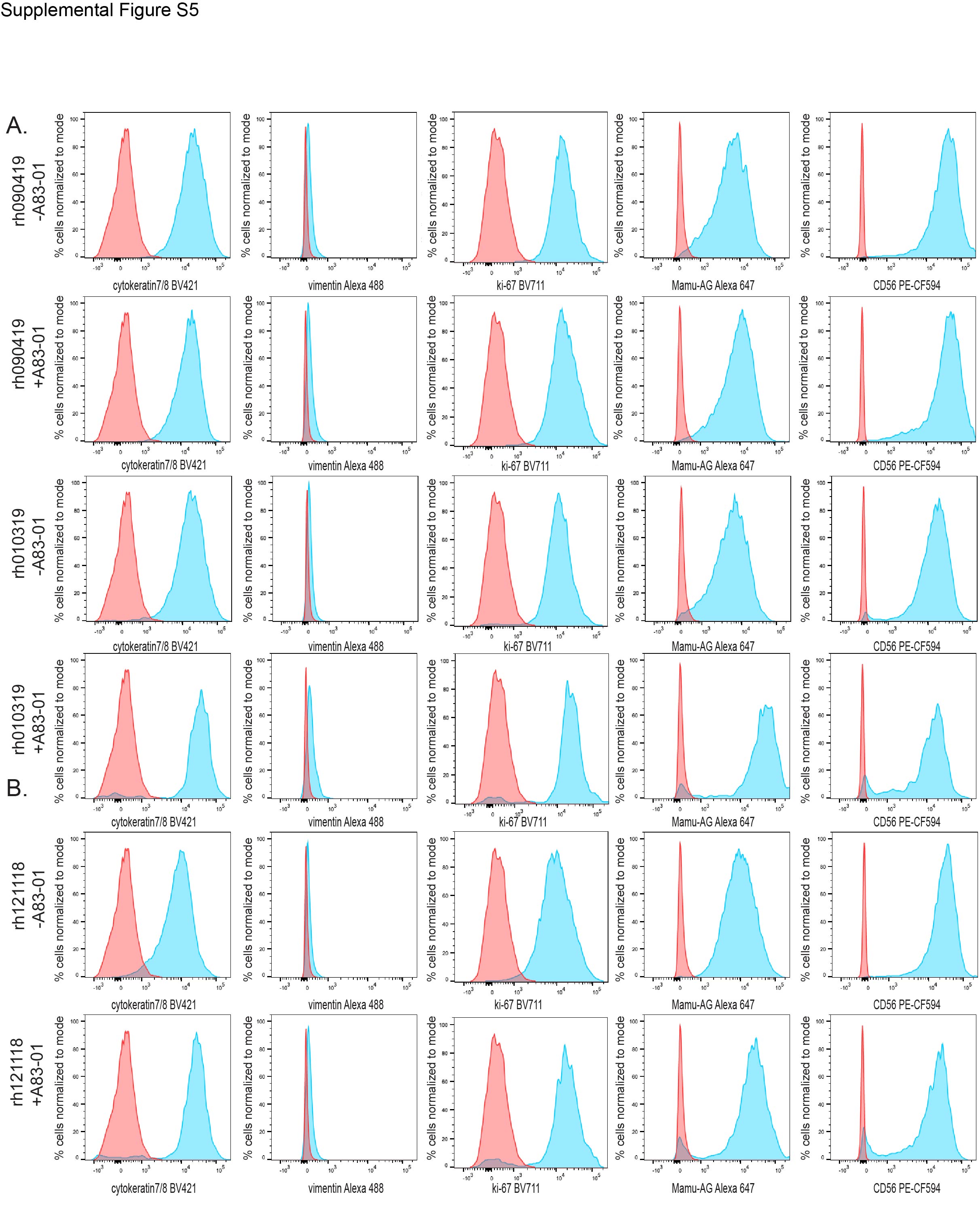
